# Barcoding of episodic memories in the hippocampus of a food-caching bird

**DOI:** 10.1101/2023.05.27.542597

**Authors:** Selmaan N. Chettih, Emily L. Mackevicius, Stephanie Hale, Dmitriy Aronov

## Abstract

Episodic memory, or memory of experienced events, is a critical function of the hippocampus^1–3^. It is therefore important to understand how hippocampal activity represents specific events in an animal’s life. We addressed this question in chickadees – specialist food-caching birds that hide food at scattered locations and use memory to find their caches later in time^4, 5^. We performed high-density neural recordings in the hippocampus of chickadees as they cached and retrieved seeds in a laboratory arena. We found that each caching event was represented by a burst of firing in a unique set of hippocampal neurons. These ‘barcode-like’ patterns of activity were sparse (<10% of neurons active), uncorrelated even for immediately adjacent caches, and different even for separate caches at the same location. The barcode representing a specific caching event was transiently reactivated whenever a bird later interacted with the same cache – for example, to retrieve food. Barcodes co-occurred with conventional place cell activity^6, 7^, as well as location-independent responses to cached seeds. We propose that barcodes are signatures of episodic memories evoked during memory recall. These patterns assign a unique identifier to each event and may be a mechanism for rapid formation and storage of many non-interfering memories.

## Introduction

The hippocampus is critical for acquiring memories across a range of timescales. Some memories form “statistically” via repeated experiences or exploration of an environment^8, 9^. Other memories form in a one-shot “episodic” manner, by an animal experiencing a single event^2, 3, 10^. Many studies have associated hippocampal activity with statistical types of learning. For example, activity of place cells reflects gradually acquired knowledge about the rewards and the geometric structure of an environment^11–16^. Much less is known about how hippocampal activity relates to episodic memory. Studies have shown that rapid, episodic-like changes to place cell activity are possible in certain situations^17–21^. Other work has proposed mechanisms beyond changes to place cells in the formation of an episodic memory^22–24^. To address this issue, it is crucial to understand how hippocampal activity encodes an individual memorable event.

Food-caching birds provide an excellent opportunity to study episodic memory^10, 25^. These birds specialize at hiding food in many concealed locations and using memory to retrieve their caches later in time^4, 26, 27^. Each caching event is a brief, well-defined episode that creates a new memory. Accurate retrieval of food caches requires the hippocampus^5, 28^, which is homologous between birds and mammals^29, 30^. Recent studies have discovered abundant place cells in birds^7, 31, 32^, suggesting that some mechanisms of hippocampal function are likely shared across vertebrate species.

We set out to record hippocampal activity in a food-caching bird, the black-capped chickadee. We engineered an experimental setup for high-density neuronal recordings in chickadees as they performed large numbers of caches, retrievals, and investigations of cache sites. We examined how the hippocampus represented individual caching events, and how this episodic encoding related to the conventional coding of place.

### Neural recordings during food caching

Chickadees form new memories during well-defined food-caching events. Our first goal was to characterize hippocampal activity during these episodes. We adapted a previously developed food-caching setup^27^, optimizing it to obtain exceptionally large numbers of caching events. The setup was an arena containing 128 caching sites concealed by cover flaps (Fig. 1a). Sunflower seeds were provided to chickadees by motorized feeders that opened only for brief periods during a session. Because caching is motivated by the instability of food supply^33^, this schedule encouraged chickadees to cache seeds whenever feeders were opened (Supplementary Video 1). When feeders were closed, chickadees spent most of the time hopping around the arena and sometimes “checking” sites by opening cover flaps. They often retrieved their caches, eating some of them and recaching others at different locations. Birds also checked sites without retrieving seeds, implying that these checks were not simply failed retrieval attempts.

**Figure 1.**
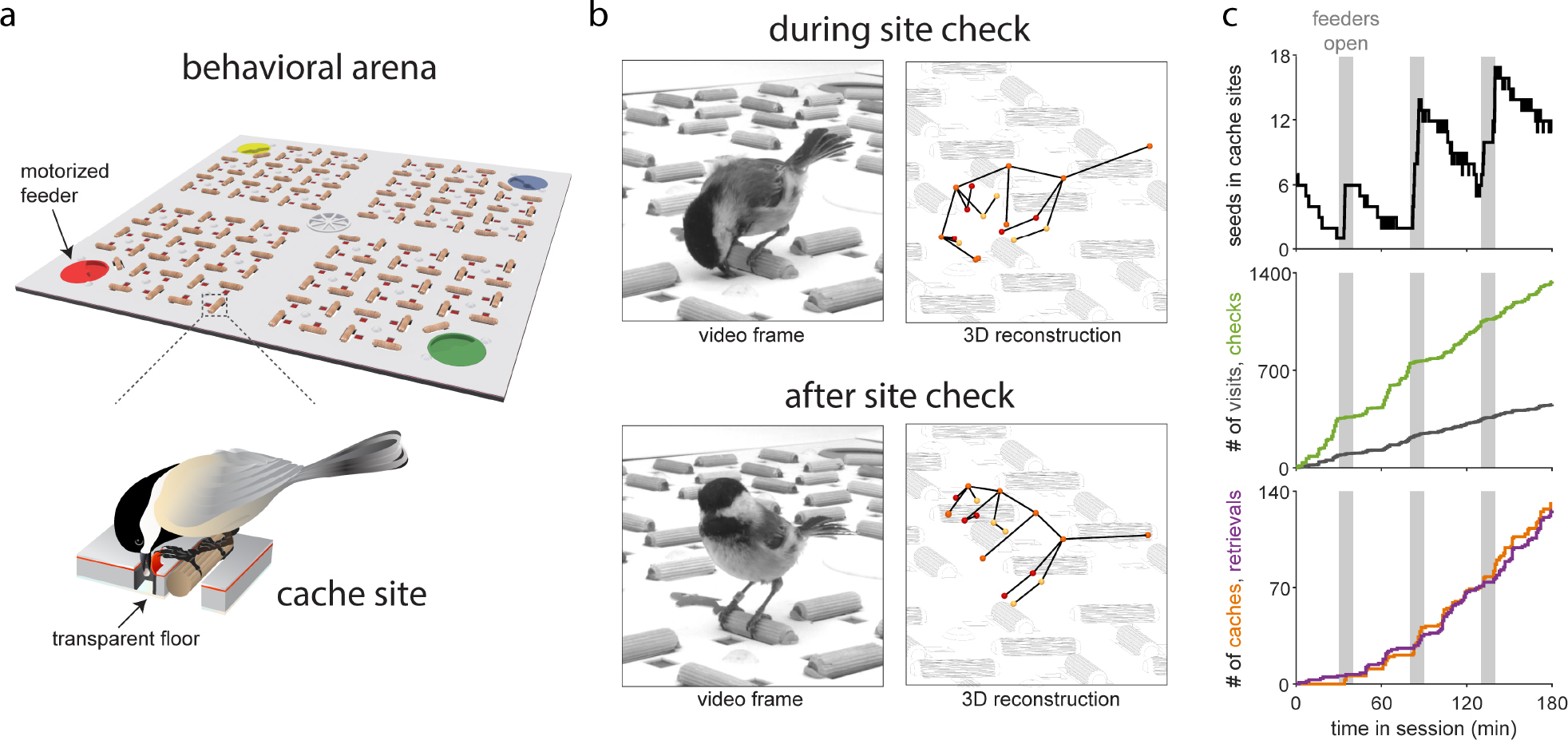
Behavioral setup for food-caching chickadees. **a)** Schematic of a 76 × 76 cm behavioral arena containing 128 cache sites with 5.3 cm minimum spacing. Each site consists of a perch and a cavity in the floor covered by a rubber flap. Chickadee lifts the cover to interact with a site. Site contents are not visible from above once the cover snaps closed, but are camera-monitored from below through a transparent floor. **b)** Left: example frames from a single video camera. Right: 3D reconstruction using six cameras, showing 18 tracked keypoints registered to the model of the arena. Keypoints are colored yellow, orange, and red for points on the bird’s left, midline, and right. **c)** Example behavioral session with three periods during which feeders were open. Chickadees cached new seeds during feeder-open periods. During feeder-closed periods they retrieved seeds and recached many of them, thus increasing the cumulative number of caches and retrievals throughout the entire session. Chickadees also “visited” sites (perch landing without interacting with the site) and “checked” sites (opening cover flap without caching or retrieving). Note that seven sites were baited at the start of the session.

We used high-resolution cameras to record behavioral videos from six vantage points in the arena. We also developed algorithms for mm-precision, 3D postural tracking of points on the bird’s body in these videos (Fig. 1b, Supplementary Video 2). An additional camera was used to detect the contents of all the cache sites through a transparent bottom layer of the arena. Automated tracking allowed us to record the chickadee’s location, to detect when its beak made contact with a site, and to determine whether seeds were placed or removed at a site. We detected four types of events: caches, retrievals, checks (opening the cover flap without caching or retrieving), and visits (landing at a site without touching the cover flap). In a typical session, chickadees made ∼100 caches and retrievals (Fig. 1c, 68-149 caches, 66-146 retrievals, 25-75% percentile, n=54 sessions). They made even more visits and checks.

Chickadees are small (∼10 g) and make dexterous head movements to cache food. To record neural activity during this behavior, we engineered a very lightweight (∼1.2 g), miniature microdrive assembly compatible with silicon probes. We also developed a “semi-acute” preparation that involved fully retracting probes from the brain between recording sessions. This procedure prevented the typical deterioration of neural signals over time and allowed us to record stable numbers of units, typically for >1 month. We recorded in the anterior hippocampus, which has been shown in other avian species to contain abundant place cells^7, 31, 32^. As in other behavioral tasks and species, the hippocampus exhibited spatially-modulated activity during site visits in our arena. Using standard criteria, 56% of the neurons were classified as place cells (2462/4366 units in 5 chickadees).

### Sparse hippocampal cache responses

Compared to visits, activity during caching events was strikingly different. We first analyzed putative excitatory cells, which are identifiable in birds using firing rates and spike waveforms^7^. These cells were largely silent during caches (Figs. 2a,b). Relative to shuffled data in which caches were randomly shifted in time, 64% of cells had a significant decrease in firing rate, and only 6% had an increase (n=2528 excitatory units, p<0.05). A typical neuron produced zero spikes during most caches, and firing rate was below its session average on 93% of the caches.

**Figure 2.**
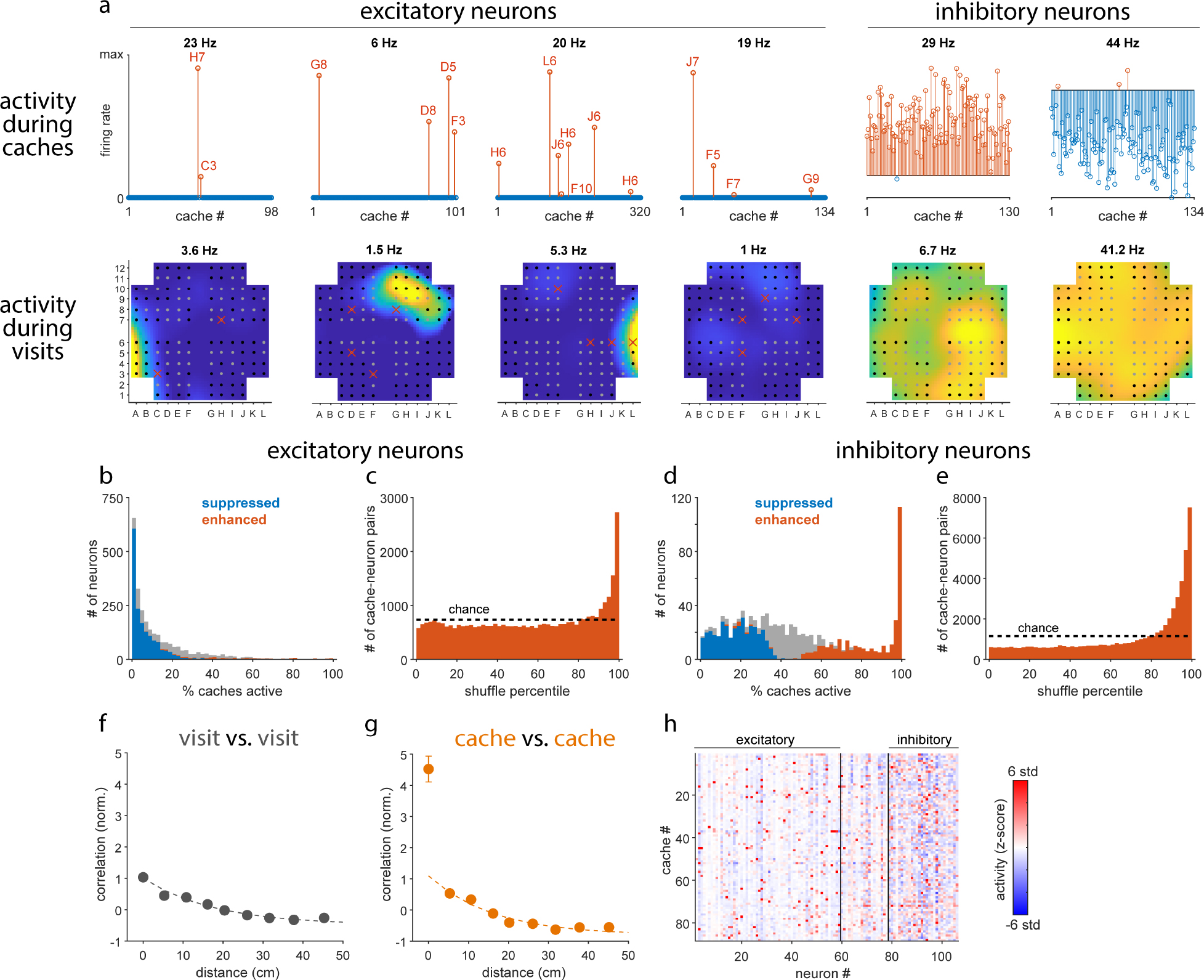
“Barcode” activity during caches. **a)** Bottom: Spatial maps for six example neurons, plotted from zero (blue) to maximum firing rate (yellow). For all cells except the fourth one, maximum is peak rate across the arena; for the fourth (“silent”) cell, peak rate was 0.15 Hz but maximum on the plot is 1 Hz. Rate was measured at sites during visits and interpolated for all other locations. Circular symbols: positions of sites. Gray circular symbols: sites that were cached into at least once in the session. Red x’s: cache locations where the neuron produced a “cache response” – i.e., was active above its mean firing rate. Top: Firing rates of the same neurons during all caches. Blue and red stems: caches with firing rate below and above the mean rate, respectively. For excitatory cells, cache responses are marked with location, to match with spatial maps below. Cache responses were sparse and not clearly related to place fields. Inhibitory cells were either non-sparsely enhanced or suppressed by caching. **b)** Fraction of caches during which an excitatory neuron responded. For most neurons, this fraction was lower than expected from shuffled data. **c)** For all cache responses, comparison of firing rate to shuffled data, considering only shuffles with above-average firing rates. Cache responses often exhibited exceptionally high rates. **d)** Same as b), for inhibitory cells. A subpopulation of these cells was strongly enhanced by caching. **e)** Same as c), for inhibitory cells. **f)** Correlation of population activity for pairs of visits. Correlation decayed gradually, indicating a smooth place code. All correlations are normalized such that the visit-visit value at 0 is 1 across the dataset. **g)** Same as f), for pairs of caches. Correlation showed a strong site-specific component of activity not explained by the place code. **h)** Activity of neurons across caches, after subtraction of the place code. We refer to this activity as the “barcode”. Firing rates are z-scored relative to shuffled data. Neurons were conservatively classified as excitatory or inhibitory, with some neurons left unclassified. Error bars in all panels: s.e.m; when not visible, they are smaller than the symbols.

We analyzed the remaining 7% of the caches, when a neuron fired above its average rate. These “cache responses” were typically scattered throughout the environment and not obviously related to the neuron’s place tuning (Fig. 2a). They often occurred well outside of a place field, and were produced at similar rates by place cells and non-place cells (Supplementary Fig. 1). Firing rates during cache responses were often exceptionally high. We compared these rates to shuffled data, counting only those shuffles that also had above-average firing rates. Expressing activity as a percentile of shuffles revealed a strong deviation from a uniform distribution, with a peak at ∼100% (Fig. 2c). In other words, excitatory neurons were mostly suppressed, but on a small fraction of caches produced some of the largest bursts of spikes ever observed in those neurons.

Different excitatory neurons were generally active on different subsets of caches, forming sparse patterns of population activity. Such a high degree of sparseness could be driven by inhibitory cells^34^. To evaluate this, we analyzed putative inhibitory cells during caching. We found that the effect on their firing was mixed, with 33% of neurons suppressed and 36% enhanced relative to shuffled data (n=1031 units, p<0.05; Fig. 2a,d). However, the strength of this effect was asymmetric. Enhanced neurons included extremely active cells that fired above their average rates on nearly every cache (>95%). These cells appeared to be a distinct subpopulation containing 14% of inhibitory units (Fig. 2d). Repeating our percentile analysis for inhibitory cells showed that during caching they also produced some of the strongest activity ever observed in these cells (Fig. 2e).

Together, our results show that caching engages a distinct state of hippocampal activity. In this state, a subpopulation of inhibitory cells is enhanced, and excitatory cells produce a very sparse pattern of firing across the population.

### Cache representation by neural barcodes

There are two major questions about the nature of the cache responses we observed. First, are they consistent for specific locations? In other words, do these responses repeat if a chickadee caches multiple times into the same site? Second, what is the relationship of these cache responses and the activity of place cells? To address these questions, we defined a population vector of activity for each caching event. We then measured the correlation of these vectors for pairs of caches at the same site and at different sites. We compared these “cache-cache” correlations to those measured for visits (“visit-visit” correlations).

Visit-visit correlation decayed roughly exponentially with distance between sites (Fig. 2f). This is expected from the activity of place cells. Because a typical place field has some spatial extent, correlation of activity is high for nearby sites, but lower for more distant sites. Similarly to visits, cache-cache correlation also showed a gradual decay at nonzero distances, indicating some smooth spatial tuning during caches. Indeed, we found that place cells continued being active during caching (Supplementary Fig. 1, considered in more detail below).

In contrast to visits, cache-cache correlation sharply deviated from a smooth function at zero distance (Fig. 2g). Caches at the same site were correlated with values ∼4.5x greater than visits at the same site. (From here on, we normalize correlation values by the visit-visit correlation at zero distance; i.e., a correlation value of 1 is expected for a pair of visits to the same site). This high correlation shows that activity was consistent across multiple caches into the same site. Signs of this can be seen in the activity of individual neurons. For example, the third cell in Fig. 2a repeatedly fired during caches into the same sites (H6 and J6).

Notably, correlation was enhanced for caches only at the same site. Cache responses even at adjacent sites just ∼5 cm apart were much lower and consistent with the presence of place cell activity during caching. This site specificity was also evident in the activity of individual neurons, which showed spatially punctate cache responses confined to individual sites (Fig. 2a). These punctate patterns were markedly different from place fields, which typically extended over multiple nearby sites.

Our observations suggest the presence of two patterns of population activity in the hippocampus. The first is the conventional, spatially smooth “place code”. This pattern is similar for nearby sites and is engaged during both visits and caches. The second pattern is highly specific to an individual site, dissimilar even between adjacent sites, and engaged only during caches. We call this site-specific activity a “barcode” – a pattern of firing in a subset of neurons against a background of suppression in the remaining neurons. A barcode is a unique representation of a cache site. It occurs transiently, only when the bird is caching a seed, but not when it simply visits the same site.

Because cache responses combined both place and barcode activity, we developed a procedure to separate these two components. To estimate the place code at a particular site, we temporarily left that site out and spatially interpolated activity recorded at other sites. Repeating this procedure for all sites produced a spatially smooth function over the environment – i.e., the place code. To estimate the barcode, we subtracted the place code from activity recorded during caches. Fig. 2h shows a matrix of residuals after this subtraction. The population vector represented by each row of this matrix is what we call the barcode.

### Reactivation of barcodes

What is the purpose of site-specific barcodes? An intriguing hypothesis is that a barcode represents a memory formed by caching at a particular site. If a barcode represents a memory, one might expect it to reactivate during other behaviorally relevant times. A reasonable starting point is to compare caches with other events at the same site. We therefore considered retrievals, checks, and visits.

We found a strong correlation of activity between caches and retrievals at the same site (Fig. 3a). This correlation began increasing ∼250 ms prior to the bird’s beak touching the site and continued for the duration of the two events (median 1.2 s for caches and 1.5 s for retrievals). There was a similar reactivation between caches and checks (Fig. 3b). In this case, reactivation was even more brief, likely reflecting the shorter duration of the checks (<0.2 s). In contrast to retrievals and checks, there was a much weaker correlation between caches and visits (Fig. 3c). This weak but positive correlation is expected from place coding during both caches and visits. These analyses show that cache-related activity was indeed reactivated, specifically during retrievals and checks – i.e., those events when the bird accessed the contents of the site. Reactivation was transient and precisely aligned to behavior.

**Figure 3.**
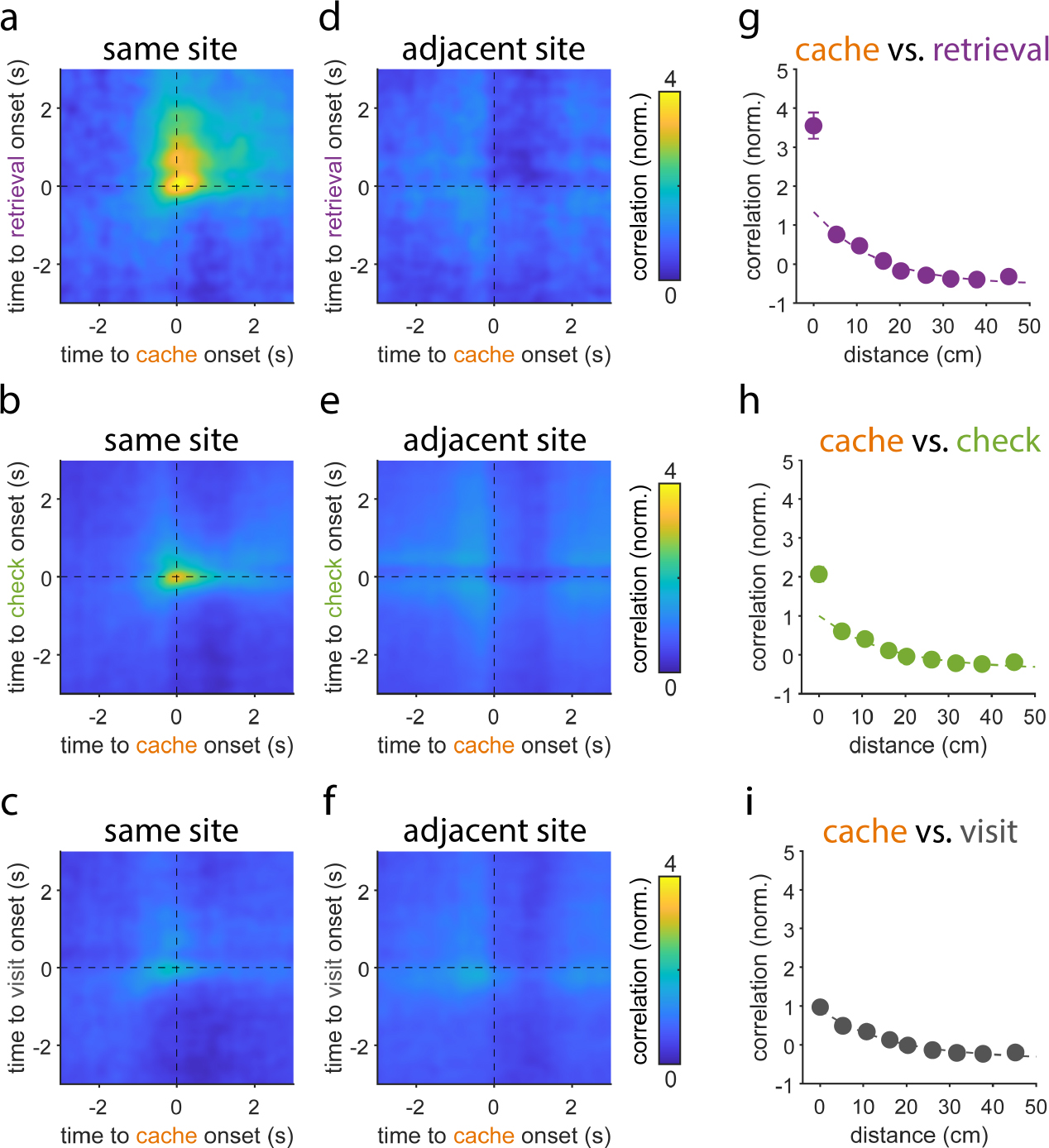
Reactivation of barcodes during site interactions. **a)** Correlation of activity during caches with activity during retrievals at the same site, averaged across all cache-retrieval pairs. Zero is the time of the bird’s beak making contact with the site cover. Population activity vectors were smoothed with a Gaussian window (σ = 100 ms). Activity was transiently reactivated between the two events. **b)** Same as a), but for cache vs. check comparison. **c)** Same as a), but for cache vs. visit comparison. Correlation was weaker and consistent with reactivation of the place code. **d-f)** same as a-c), but comparing caches with events at adjacent sites (5 cm away). Reactivation was weak in all cases. **g-i)** Correlation of activity during caches with activity during other events, as a function of distance between sites. There was a gradual decay in all cases, demonstrating the presence of a smooth place code. During retrievals and checks, there was also a reactivation of the site-specific barcode. Error bars in all panels: s.e.m; when not visible, they are smaller than the symbols.

Our analysis also showed that reactivation was highly site-specific. Strong correlation was not observed when we compared caches at one site to events even at adjacent sites (∼5 cm away; Figs. 3d-f). For all types of events, correlation with caches decreased roughly exponentially as a function of distance (Figs. 3g-i). For retrievals and checks, however, correlation also showed a narrow peak at zero distance. This suggests that the smooth place code was active during all events, but there was an additional reactivation of the barcode during retrievals and checks. We confirmed this by isolating the barcode from cache-related activity using the procedure described above, and by correlating neural activity to the barcode. Correlation to the barcode was high for retrievals and checks at the same site (3.0±0.25 and 1.6±0.14, mean±sem, n=54 sessions) and lower for visits (0.55±0.10). Correlation between the barcode at one site and all events at adjacent sites was low (0.25±.06, 0.17±0.02, and 0.07±0.02). Finally, we adapted this analysis to determine which individual neurons contributed to barcode reactivation. We found significant barcode reactivation in 29% of excitatory and 30% of inhibitory neurons, with no relationship between a neuron’s place and barcode activity (Supplementary Fig. 2).

### Memory drives changes to barcodes

Why are barcodes reactivated across different events? Caches, retrievals, and checks are fairly similar; for example, they involve similar motor actions by the bird. Therefore, one possibility is that the barcode, and its reactivation, represent the conjunction of a particular action with a particular location. An alternative possibility is that a barcode represents a memory of an event, and that this memory is recalled at behaviorally relevant time points. If this is the case, we might expect the barcode to contain additional information that is unique to a specific caching event.

To test these possibilities, we first asked whether barcodes were different across caching events at the same site. Our previous analysis showed that barcodes at the same site were correlated. However, this correlation does not exclude the possibility of meaningful differences between the barcodes as well. Indeed, we found that there were systematic differences between barcodes. Barcode-barcode correlation was high for pairs of consecutive caches at the same site, but decreased for pairs separated by intervening caches (Fig. 4a). This effect could result from the animal’s experiences (i.e. intervening caches), or merely reflect drift in neural activity over time. To test this, we analyzed correlations for consecutive barcode pairs as a function of the time interval between caches (Fig. 4b). There was a short-latency effect for pairs separated by less than 5 min. However, at longer intervals correlations approached an asymptote at a value of ∼5, much higher than correlations between barcodes separated by intervening caches. This suggests that the decrease in barcode-barcode correlation is due to the animal’s experience, rather than elapsed time. We obtained similar results using a linear mixed-effects model which accounted for multiple variables simultaneously (Supplementary Fig. 3a). In summary, barcodes were not constant at a given location, but were distinct for each caching event.

**Figure 4.**
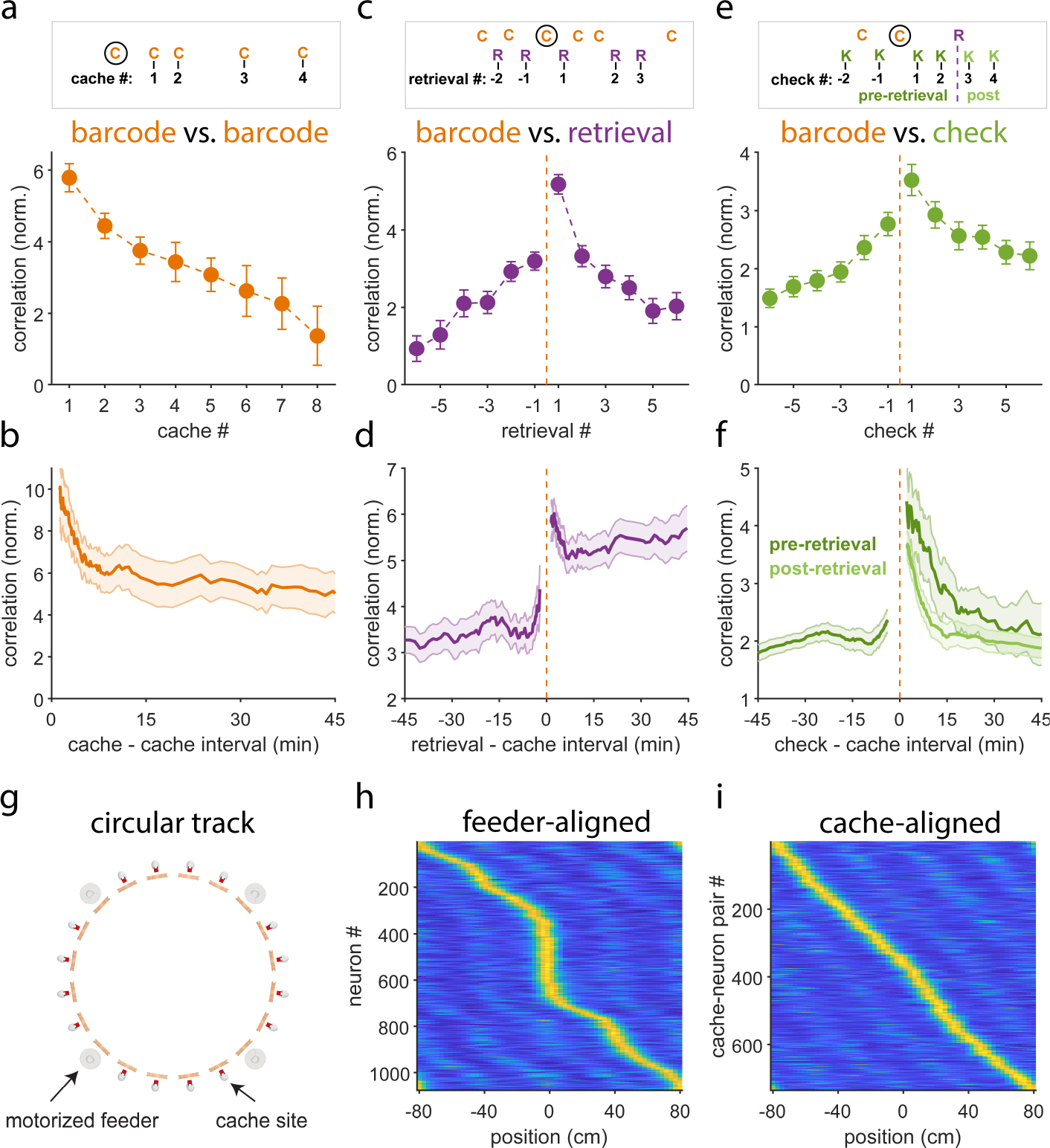
Representation of specific caching episodes by barcodes. **a)** Top: schematic of the analysis. Barcodes during caches (“C”) that occurred at the same site but at different times are compared to each other. Cache # indicates the separation of a cache from the one indicated by the black circle: e.g., 1 indicates two consecutive caches, 2 indicates two caches separated by one intervening cache, etc. Bottom: average correlation of barcodes as a function of cache #. Increasingly separated caches had increasingly different barcodes. **b)** Correlation of two consecutive caches as a function of their separation in time. The trace asymptoted at a high value, indicating that barcodes became less similar not due to elapsed time, but due to intervening caching events. **c)** Top: schematic of the analysis, with retrievals (“R”) assigned a retrieval #. Retrieval #1 is “matched” to the cache indicated by the black circle. **d)** Correlation of the barcode to the retrieval, as a function of retrieval #. Matched retrieval (#1) had the strongest correlation to the barcode. **e**) Correlation of the barcode to retrievals immediately preceding and following a cache (#’s -1 and 1) as a function their separation in time from the cache. Barcode was most strongly correlated to the subsequent retrieval, and this effect was stable for long time intervals. **e)** Same as c), but for checks (“K”). Barcode was most strongly correlated to the check that immediately followed the cache. **f)** Same as e), but for checks. Checks are analyzed separately depending on whether they occurred before or after the retrieval that followed the cache. Barcode reactivation during checks was strong immediately after a cache, but did not persist at long time intervals. The decay in reactivation strength was even faster after the seed was retrieved. **g)** Schematic of the setup for analyzing the effect of caching on place cells. Arena is similar to the one shown in Fig. 1, but has cache sites arranged along a circular track of 160 cm circumference. **h)** Alignment of activity to the rewarded feeder, which was changed on a session-by-session basis. Each row is a neuron’s spatial tuning curve around the circular track, unwrapped. Rate is normalized from 0 (blue) to the peak (yellow) for each cell. For clarity, only neurons that had a peak firing rate exceeding 3 standard deviations of the spatial tuning curve are shown. Place fields were overabundant near rewarded feeder. **i)** Alignment of activity to sites containing cached seeds. Place fields were not overabundant near caches. Error bars in all panels: s.e.m.

This analysis suggests that a barcode does not simply represent the act of caching at a particular site, but encodes a specific caching episode. To test this idea, we asked whether reactivations of barcodes were also distinct for different caching events. At each site, a chickadee produced some sequence of caches and retrievals. In this sequence, a “matched pair” was a cache paired with the first retrieval that followed it. We reasoned that recall of a memory formed during caching should be most likely during its matched retrieval. We found that barcode-retrieval correlation was indeed the strongest for matched pairs (Fig. 4c). Correlation was weaker for all later retrievals, as well as for retrievals that preceded the cache. In other words, activity during retrievals reactivated barcodes of specific, matched caching events.

Can this effect be explained by temporal proximity of the cache and the following retrieval? Across the dataset, matched pairs of caches and retrievals were separated by a wide range of time lags, from seconds to tens of minutes. We analyzed the first retrieval that followed a cache and the last one that preceded it, as a function of the time lag (Fig. 4d, Supplementary Fig. 3b). Again, there was some short-latency effect in the data: caches and retrievals were more correlated when separated by less than ∼5 min. At all lags, however, reactivation was stronger for matched pairs of caches and retrievals. The barcode was more strongly correlated to the following retrieval than to the preceding retrieval – even at lags of 45 min.

In summary, the hippocampus produced a distinct barcode during each caching event. This event-specific barcode was reactivated during a subsequent retrieval from the same site. Reactivation occurred even after long delays. These results suggest that the barcode represents a specific episodic experience.

We repeated these analyses for checks. Barcode-check correlation was also stronger for checks that followed a caching event (Fig. 4e). However, reactivation during checks was not as persistent as during retrievals. Barcode-check correlation decayed back to baseline with a timescale of 13.8 min (Fig. 4f, Supplementary Fig. 3c). Note that this does not imply a disappearance of memory; in fact, a retrieval after this time would successfully reactivate the barcode. Rather, reactivation was context-dependent: sometime after caching, it stopped occurring during checks and only occurred during retrievals. After retrieval, the barcode-check correlation decayed to baseline even faster, with a timescale of 5.0 min (Fig. 4f, Supplementary Fig. 3d). In other words, once a cache was retrieved, the hippocampus quickly stopped reactivating the corresponding barcode.

### Place cells are unchanged by caching

Our results show that cache memory is represented by transient hippocampal barcode activity. Are caches also represented by the place code? Place maps are known to change in experience-dependent ways across a number of experimental conditions^11–16^. We thus asked whether place cell firing was also different before and after caching. This was challenging in our 2D arena due to limits in sampling, since obtaining 2D maps requires sufficient coverage of the environment both before and after a cache. We therefore designed a 1D version of the arena, in which chickadees moved and cached around a circular track (Fig. 4g). The advantage of this track is that behavior was more repeatable: birds typically visited the same site many times before and after a cache, allowing a comparison of activity across many trials. We recorded in 7 additional chickadees on this track.

In rodents, place fields can shift with experience and tend to be overabundant around reward locations^12, 15, 35^. In our experiments, the location of a single rewarded feeder was changed from session to session. Consistent with rodent data, place fields were concentrated around the rewarded feeder in chickadees (p<0.001 KS test for uniform distribution, Fig. 4h). In contrast, we did not find a similar redistribution of fields around caches (Fig. 4i). Aligned to cached seeds, place field locations did not significantly deviate from a uniform distribution (p=0.12 comparison to uniform distribution; p<0.001 comparison to deviation from uniformity around rewarded feeder).

We also looked for other types of changes to the place map. Across published experiments, place fields have been observed fully remapping^17, 36, 37^, changing their firing rates^37–39^, changing their widths^40, 41^, changing their shapes^40, 42^, or shifting their locations^12, 15, 35^. We did not observe any of these changes in the food-caching behavior (Supplementary Fig. 4). Comparison of spatial activity before and after a cache did not show differences greater than expected from randomly shuffling cache times. Although we cannot rule out more subtle effects, cache memory seems to be mainly represented in the transient barcode activity rather than in the place code.

### Responses to seeds in the hippocampus

So far we have focused on the patterns of activity that are spatially selective. These include both place codes and barcodes. However, neurons also had non-spatial changes to their firing. As we showed earlier, many cells were either suppressed or enhanced on average across all caches, regardless of their location (Fig. 2a). What causes these changes to the average firing rate? We examined whether this cache modulation might itself be encoding a non-spatial aspect of the chickadee’s experience.

An important variable for the chickadee is the presence or absence of a seed in a site. Checks provide an opportunity to study this variable, because chickadees check occupied sites that contain a cache, as well as empty sites. Checks were the most frequent action performed by chickadees, and were very brief and stereotyped (150-200 ms). We found that many neurons had different firing rates between occupied and empty checks, regardless of location (Fig. 5a). These differences were especially prominent in inhibitory cells: they were significant in 36% of inhibitory and only 4% of the excitatory units (p<0.01). We defined a “seed vector” as the difference in population activity between occupied and empty checks. We then measured the strength of the “seed response” by projecting population activity onto this vector. The response was cross-validated by holding out one check at a time when computing the seed vector. Seed responses diverged between occupied and empty checks ∼100 ms after the bird lifted the cover flap and peaked just after check offset (Fig. 5b). The seed response was absent when the chickadee visited a site without checking (Figs. 5c,d).

**Figure 5.**
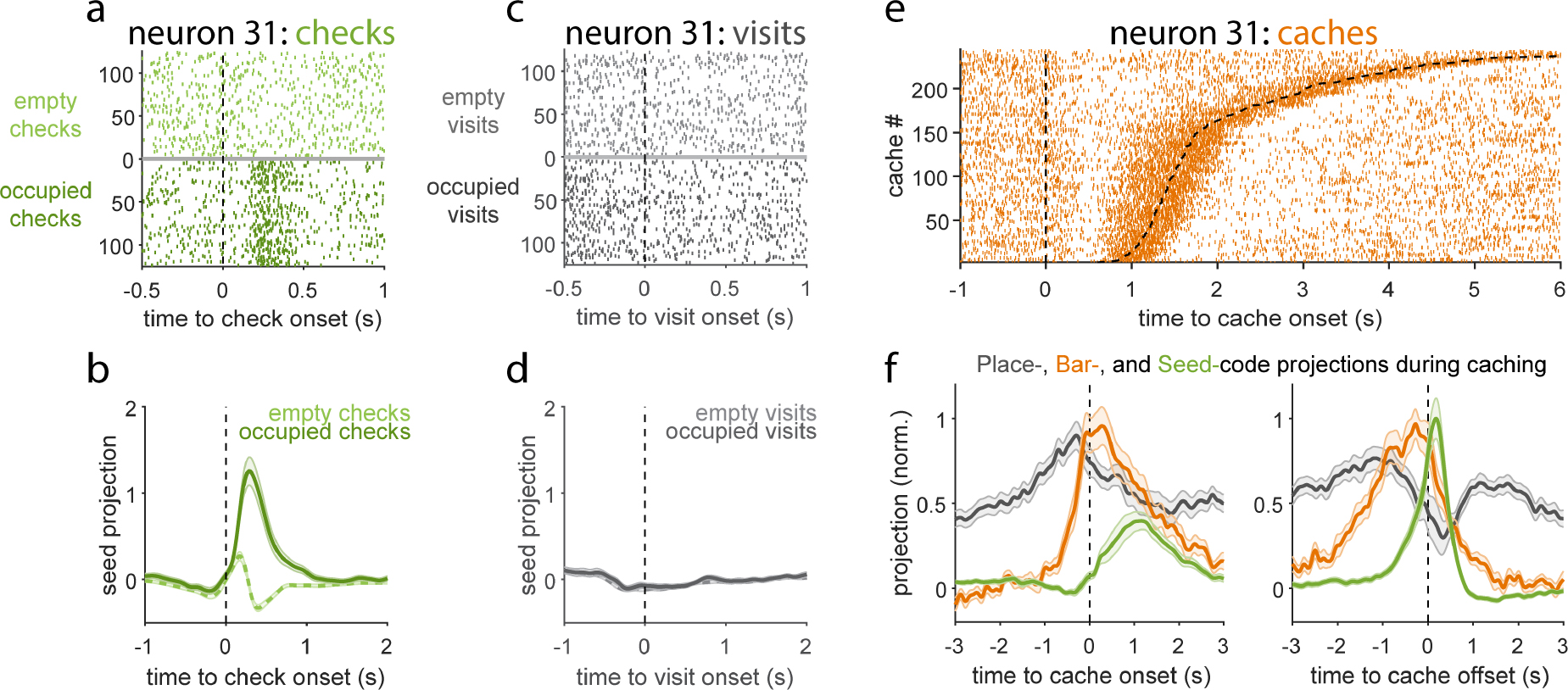
Coordination of different neural codes during caching. **a)** Raster plot of activity during checks. Zero is the time of the bird’s beak making contact with the site cover. For display purposes, the same number of empty and occupied checks was selected randomly from the session. Neuron showed a response to the presence of a seed in the site. **b)** Average projection of neural activity onto the “seed vector”, defined as the difference in population activity between occupied and empty checks. **c)** Activity of the same neuron as in a), but during visits. Response to the cached seed is absent. **d)** Same as b), but for visits. **e)** Activity of the same neuron as in a) and c), but during caches. Caches are ordered by duration, with both onset and offset shown (dashed lines). Neuron responded around cache offset. **f)** Average projection of neural activity during caches onto vectors defined by the place code, the barcode, and the seed code. Data were smoothed with a Gaussian window (σ = 47 ms). Activity was aligned separately to onsets and offsets of caches; median cache duration was 1.2 s. Error bars in all panels: s.e.m.

We found that the seed response was strong not only during occupied checks, but also during caches (Figs. 5e,f). Interestingly, this response was tightly locked to the offset of the cache – i.e., the moment in time when the chickadee left a seed in the site. This offset alignment was evident in the population seed response and in the activity of individual neurons. The seed response was not as strong prior to this moment, even though the chickadee had first carried the seed to the cache site, and then interacted with the cache site for over 1 s. This pattern of activity thus seemed to occur when the bird attended to a seed that was in a cache site, both during checks and caches. We wondered how this time course compared with other patterns of activity.

We projected population activity onto cross-validated place code and barcode vectors for each site. We found that all three responses followed different time courses (Fig. 5f). The place code peaked prior to the onset of the cache, roughly when the chickadee arrived at the site. The barcode started increasing prior to the onset and peaked during the cache itself. Finally, the seed response peaked at the offset of the cache. Place, barcode and seed projections showed similar temporal differences when computed for visits, checks, and retrievals, although the magnitude of barcode and seed projections were less than during caches (Supplementary Fig. 5). These results show a complex sequence of hippocampal responses during caching, synchronized with behavior on a sub-second timescale.

## Discussion

Our recordings uncover sparse patterns of hippocampal activity (“barcodes”) that uniquely represent food-caching events. These patterns reactivate when a chickadee checks or retrieves a cache, even long after caching. We call these responses “barcodes”, since they provide a seemingly random index for each event, although the metaphor should not be taken too literally. For example, hippocampal activity was sparse and continuous-valued, whereas literal barcodes are typically dense and binary. Barcodes coexist in hippocampal activity with the conventional place code, but are notably different. Whereas the place code is persistently active during movement, barcodes activate transiently, only for the brief time that it takes a chickadee to interact with a cache site. Whereas the place code is spatially smooth across the environment, barcodes are uncorrelated even for adjacent sites. Finally, barcodes are not stable representations of location or the act of caching: they are different even across caching events at the same site. These properties are consistent with a barcode being a unique signature of a specific cache memory, evoked when the chickadee recalls that memory at a behaviorally relevant time.

A growing body of work has shown that experience can also modify the firing of place cells. For example, place maps of different environments gradually diverge as an animal spends time exploring these environments^14^. Place fields also reorganize to over-represent rewards and other salient locations^12, 15, 35, 43^. Changes to place fields can even be sudden, suggesting a capacity for rapid learning^17–21^. In contrast to these studies, we have not observed significant changes to the place code resulting from food-caching events. This finding implies a dissociation of hippocampal mechanisms for different types of memory. Acquired knowledge about consistent features of an environment may be represented by the place code. On the other hand, specific events within that environment appear to be represented by barcodes.

Our analysis shows a precise temporal coordination of barcodes with activity representing location (the place code) and cached food (the seed code). During a caching event, all these codes occur within a window of ∼1 s, compatible with known mechanisms of fast hippocampal plasticity^20, 21, 44^. It is conceivable that a similar mechanism in birds synaptically links neurons that participate in these codes. One speculative idea is that memory formation involves connecting barcode neurons with both place cells and seed cells, in effect making them a link between representations of place and food. In this model, barcodes are unique identifiers of individual memories. They bind together different components of a memory, but prevent interference that would arise if, for example, many nearby place cells were directly linked to the same neuron representing a sunflower seed. As a result of this binding, partial reactivation of place or seed inputs – e.g. when a bird is searching for a nearby cache – could reactivate the barcode. This hypothetical mechanism would permit a bird to recall previous cache memories in a contextually appropriate manner.

Our work parallels ideas about episodic memory from other systems. In the human hippocampus, activity patterns that occur during memory formation are subsequently reactivated during episodic recall^45–48^. In animal models, the concept of an “engram” is often used to describe cells that represent a specific memory^49, 50^. There is an ongoing inquiry into the nature of engrams. In the traditional definition, an engram includes all cells that are reactivated between memory formation and recall^23, 49, 50^. In our data, these cells would include not only the barcode, but also place cells and seed cells. More recent studies have proposed treating cells that were already active before a memory (such as place cells) separately from those that become newly active during memory formation^22, 23^. These newly active cells appear to be a small fraction of the population and might be active only transiently during recall of a specific memory. Our findings about barcodes are consistent with this concept.

Our results make it tempting to conclude that barcode reactivation is a mechanism of memory recall. We have not yet shown this to be generally the case. Reactivation during checks and retrievals happens when the chickadee is already at a cache site, about to open the cover flap. In contrast, memory recall should also happen in advance, at a time when a chickadee makes a choice about where to go next. This is especially important for an animal navigating through a large, naturalistic environment that is very sparsely populated by food caches. We predict that in complex environments, barcodes activate at decision points, and that this activation influences subsequent behavior of the bird. This prediction remains to be tested in future work.

## Contributions

S.N.C., E.L.M., and D.A. conceived the study. S.N.C. performed and analyzed behavioral and electrophysiology experiments, and developed methods for 3D tracking. E.L.M. performed and analyzed calcium imaging experiments. S.H. acquired animals and assisted with animal care, data analysis, and experiments. S.N.C. and D.A. wrote the manuscript. D.A. supervised the project.

## Acknowledgments

We thank D. Biderman and L. Paninski for help with pose tracking methods, H. Payne for contributions to developing electrophysiology techniques, L. Abbott and A. Williams for conceptual advice and discussions, the Black Rock Forest Consortium, J. Scribner and the Hickory Hill Farm, and T. Green for help with field work, and members of the Aronov laboratory for comments on various portions of the manuscript. Illustrations in Fig. 1a are by J. Kuhl. This research was supported by the Beckman Foundation Young Investigator Award, the New York Stem Cell Foundation – Robertson Neuroscience Investigator Award, NIH Director’s New Innovator Award (DP2-AG071918), NIH BRAIN Initiative Postdoctoral Fellowship (S.N.C, 5F32MH123015), Simons Society of Fellows (E.L.M.), and NIH Pathway to Independence Award (E.L.M, 1K99NS121256).

## METHODS

### Experimental subjects

All animal procedures were carried out following US National Institutes of Health guidelines, and approved by the Columbia University Institutional Animal Care and Use Committee. Five black-capped chickadees (three male, two female) were used for electrophysiological experiments in the 2D arena, and seven chickadees (one male, four female, two undetermined) were used for calcium imaging experiments on the circular track. Subjects were collected from multiple sites in New York State using Federal and State scientific collection licenses. Chickadees are not clearly sexually dimorphic, and all experiments were performed blindly to sex. Subject age at collection time was not precisely determined. Birds were collected between October and February, and data were acquired between three months and one year after collection. Birds were housed in groups of 1-3 on a “winter” light cycle (9:15 light:dark hours) before experiments began.

### 2D arena design

The arena for 2D behavior and electrophysiology was adapted from a previously published design^1^. It was designed in Autodesk Inventor and constructed from five layers of laser-cut material. The top layer was a 1.5 mm thick matte white acrylic sheet. The 2^nd^ layer was 1/32” thick 60A durometer synthetic rubber sheet. The 3^rd^ layer was a 1 mm thick clear acrylic sheet. The 4^th^ layer was a 5.6 mm thick black acrylic sheet. The bottom, 5^th^ layer was a clear 1/8” thick acrylic sheet. Wooden dowels 3/8” in diameter and 1.5” long were used for perches. Perches were aligned with slots laser-cut through all layers of the arena, and were secured using Loctite Fun-Tak. Cache sites were formed by a 0.3 × 0.25” hole cut into the 4^th^ arena layer, with the 5^th^ layer forming a transparent bottom. The 2^nd^ arena layer was cut to form a rubber flap which fully covered the underlying cache site, and the 1^st^ arena layer was cut to allow access to the underlying rubber flap. The 3^rd^ arena layer served as a spacer between the rubber flap and the 4^th^ arena layer, in order to allow the bird to easily grab the flap with its beak.

The arena was designed as 4 symmetric quadrants, each containing 32 cache sites with one perch per site. Sites were positioned such that the midpoints between the centers of the sites and their matched perches lay on a 6×6 rectangular grid with 5.3 cm spacing. Sites were grouped in modules of 4 sites facing each cardinal direction. Four sites at the outside corner of each quadrant were eliminated to permit room for a feeder, resulting in a total of 128 cache sites. Feeders were constructed from 3D printed material (red, blue, green, and yellow Processed Versatile Plastic, Shapeways). A shallow dish holding sunflower seeds was covered by a top piece controlled by a stepper motor, allowing automated opening and closing of access to the dish beneath. The stepper motor was controlled by Arduino and a stepper motor driver. A perch was placed next to each of the feeders. Finally, a water dish was 3D printed (White Resin, Formlabs 3 printer) and inserted into a circular cutout in the center of the arena.

The entire arena was mounted on a custom-constructed aluminum frame. The frame also supported lighting and video cameras described below. A 6” border constructed of matte white acrylic surrounded the arena and was also enclosed within the frame. Arena walls were constructed from white vinyl shower curtains cut to size and secured to the external frame. A single orienting cue (11 × 8” black rectangle) was positioned in the center of one of the walls. Additional cues were 12 small stickers of varying colors and shapes placed on the floor of the arena in the space between arena quadrants.

Behavioral videos were collected using six Blackfly S cameras (BFS-U3-70S7M-C, Flir Teledyne, SONY IMX428 monochrome sensor) using wide-angle lenses (8 mm focal length, M111FM08, Tamron). Four of these cameras were positioned roughly at eye level with the chickadee in the corners of the arena. Two additional cameras were mounted on the ceiling (60 cm above the floor) at the midpoints of two opposite edges of the arena. Each of the six cameras was oriented to obtain a complete view of all the sites and feeders in the arena. We used 800 µs exposure times to minimize blur, and frame acquisition was synchronously triggered across all cameras at 60 Hz rate. We used PIMAQ software (https://github.com/jbohnslav/PIMAQ) to acquire video data. Videos were compressed online during acquisition to h.264 format using two NVidia RTX2080ti GPUs. Calibration for 3D registration of video data was performed using a laser pointer (https://github.com/JohnsonLabJanelia/laserCalib). A seventh camera was positioned beneath the arena in order to monitor the contents of cache sites. Frames of this camera were triggered synchronized with every other behavioral frame acquisition (i.e. at 30 Hz). The arena was illuminated by white LED panels (superbrightleds.com).

### Behavioral protocol

At least one week before experiments, chickadees were transferred from colony to single housing. They were provided ad-libitum Mazuri small bird diet and weighed daily. Primary wing feathers were trimmed to prevent flight and promote ground foraging in the behavioral arena. A miniaturized assembly containing 4 cache sites identical to those used in the behavioral arena was baited with sunflower seeds and installed in the bird’s home cage to permit familiarization with the cache site mechanism.

Before a chickadee woke on the morning of a behavioral experiment, all food was removed from its cage. The bird was food deprived and monitored regularly, typically for the first three hours of the day. It was then placed in the behavioral arena. The arena contained a fresh water tray and four motorized feeders containing raw, shelled sunflower seeds chopped into halves. In a typical session, 6-8 cache sites were baited before the bird entered the arena. Feeders were closed for the initial 20 min and opened for 6 min every 50 min the bird remained in the arena. The exact duration of food-deprivation, number of initially baited sites, and precise feeder schedule were adjusted on a bird- and session-dependent basis, in order to optimize the bird’s caching behavior and engagement with the arena. Sessions were typically 120-180 min long. Birds were given at least one full day of rest with an ad-libitum food supply between experimental sessions.

Prior to any surgical manipulations, we ran behavioral habituation sessions. Many chickadees performed caching behavior in the arena during their initial exposure. Some birds required 2-4 behavioral sessions before readily caching and retrieving seeds, potentially due to the stress of handling and the novel environment, or unfamiliarity with the cache site mechanism. Behavioral habituation was continued until a bird appeared to comfortably perform caching behavior throughout a full session. A fraction of birds (∼1/4) either did not exhibit motivation to cache, or did not engage with cache sites, and were excluded from further experiments. For included subjects, we excluded a small fraction of sessions where the bird performed under 30 caches during a session.

### Postural tracking

Neural networks were used to track the animal’s 3D posture during behavioral sessions. We used a custom implementation of a two-stage algorithm, inspired by the DANNCE algorithm^2^, and built using the DeepPoseKit framework^3^ in TensorFlow 2^4^. The first stage of the algorithm consisted of a Stacked DenseNet, with two stacks and a growth rate of 40, which was trained to identify the coarse location of the bird’s head, body, and tail in 4x spatially downsampled behavioral videos. The bird’s body position in all 6 camera views was then used to triangulate 3D position, and the full-resolution video from each view was resampled and cropped such that the bird was centered and at constant physical scale. The second stage of the algorithm consisted of another Stacked DenseNet (2 stacks, growth rate 40) trained to detect 18 keypoints on the bird. Keypoints were chosen as reliably visually identifiable points on the bird’s exterior that did not completely align to its underlying skeleton. These included the top and bottom tips of the beak, the top and back of its head, the centers of its back and front chest, and the base and tip of its tail. They also included the left and right eyes, lower corners of its bib marking, shoulders, ankles, and feet. The output of the second stage was the location and detection confidence of all of these 18 markers in all 6 camera views. Conversion from pixel to 3D coordinates was performed using standard triangulation techniques, for each frame utilizing all pairs of the four camera views with greatest confidence rankings for that frame, and taking the median across pairs.

Training data were prepared using Label 3D software (github.com/diegoaldarondo/Label3D). An initial set of 360 frames (or 2160 images using all 6 cameras) was manually annotated. The tracking algorithm was then run on new data, and new training data were iteratively selected using a consistency metric across views (the reprojection error) to identify postures with poor tracking performance. This procedure was continued until reprojection errors were ∼1 pixel, after labeling 586 frames (3516 images). Accuracy was judged by subjectively evaluating videos, and by comparing predictions of the algorithm with two human annotators. Tracking was approximately as consistent with either annotator as the two were with each other (∼1 mm positional difference). For analysis used in the paper, tracked coordinates were then post-processed using a Kalman filter to enforce smoothness, and to interpolate over rare, brief intervals where tracking was inconsistent across views (reprojection errors >12 pixels).

### Action identification and neural window definitions

Continuous timeseries of 3D postural tracking data were parsed into a sequence of discrete actions. We first identified two kinds of events: movement between sites (i.e. the bird’s feet landing on or leaving a perch), and interaction with a cache site (i.e. the tip of the beak coming into contact with the rubber flap covering the site).

We identified movement between sites by detecting “perch arrivals” and “perch departures”. These were defined by determining when the chickadee’s feet entered or exited a 2D bounding box surrounding each perch in the arena, excluding time points where the feet were moving rapidly (>20 cm/s). Chickadees rarely left one perch without hopping to a new perch. We identified cache site interactions by identifying periods when the bird’s beak was within a 2D bounding box surrounding each cache site, and below a height threshold of 4 mm above the arena. If multiple interactions at the same site were identified within 1 s of each other, they were merged into a single longer-duration interaction.

For the analysis in this paper, actions were further subdivided into visits, checks, caches, and retrievals. These actions occurred in variable sequences and had different durations. For analysis of neural data aligned to these actions, we defined temporal windows that minimized bleed-through between them. We thus examined all perch arrivals, and identified if the bird made any interaction with the cache site before departure. Perch arrivals with no subsequent interaction of any kind were identified as “visits” in the main text. To define the window of the visit, we started with a window of ±500 ms from perch arrival, and then further refined this window to exclude confounding events. Specifically, the visit window was adjusted to begin after the offset of any interactions at other sites occurring before a visit at this site, and the visit window was truncated early if perch departure occurred less than 500 ms after arrival. Neural data during visits were defined by averaging neural activity within this window.

Caches and retrievals were defined as any cache site interactions resulting in the addition or removal of a seed, following procedures detailed in the next methods section. Caches and retrievals were extended events, typically lasting >1 s, with a heavy-tailed duration distribution. We defined a window starting 250 ms before the onset of an interaction, and extending 250 ms after the offset of the interaction. This window was further refined to start no earlier than departure from the previous perch, and to end no later than arrival at the next perch. Finally, the window for long site interactions (>2 s, ∼5% of interactions) was adjusted to include only time periods up to 1 s after onset and up to 1 s before offset, excluding the times between. Neural data during caches and retrievals were defined by averaging neural activity within these windows.

The majority of each bird’s site interactions (∼75%) were extremely brief, and involved a stereotyped motor program lasting 150-200 ms, during which the bird lifted the rubber flap with its beak and quickly peeked at site contents. We called these actions “checks”. In order to ensure we were analyzing a highly stereotyped action, we used a Gaussian mixture model to classify postural timeseries aligned to site interaction onsets. Specifically, for all site interactions that were neither cache nor retrieval, we collected the following features derived from postural tracking: (1) height of the beak above the arena; (2) distance of the beak in the horizontal 2D plane from the center of the cache site; (3) the vertical angle of the head, determined by the vector between the midpoint of bird’s eyes and the tip of its beak; (4) the distance of the beak tip from the bird’s feet. We collected data for each feature as a 25-frame timeseries from -100 to +300 ms relative to interaction onset. We visualized the data using tSNE embeddings and manually defined the cluster of events in a dataset containing 47,913 site interactions taken from 3 sessions from each of the 5 birds used for electrophysiology. We also manually defined clusters for other less common site interaction such as long interactions, swiping the beak across the cache site, or touching cache sites which were not matched to the perch where the bird was positioned. A Gaussian mixture model was then defined by these features and manual labels, and used to classify all site manipulations in the dataset. Checks analyzed in the manuscript were defined as the 80% of site interactions (excluding those with a cache or retrieval) classified as a stereotypical short check by the Gaussian mixture model. The other site interactions were excluded from analysis. The window used to average neural data for a check was from 250 ms before onset to 250 ms after offset. This window was further refined as for caches and retrievals to exclude overlap with movement between perches.

We examined alternative criteria for defining windows for each action, including windows ranging from 250 ms to 2 s wide aligned to event onsets. Results were not qualitatively affected by the choice of window.

### Detection of caching and retrieval

In addition to tracking 3D posture, we developed neural networks for semi-automatically identifying the bird’s seed handling, i.e. caching, retrieving, and other interactions with the cache site. We used video from the camera positioned below the arena, which could view the contents of cache sites through the transparent bottom. We cropped videos into 51×51 pixel bins centered on each cache site, and trained a neural network to predict whether each site was empty or occupied by at least one seed. The network consisted of layers with: 10 7×7×1 convolutions with stride 2, 25 3×3×10 convolutions with stride 1, 50 3×3×25 convolutions with stride 1, global average pooling, 25% dropout, and a 10 unit fully connected layer before a 2 unit softmax classification output. We applied ReLU activations, batch normalization, and 2×2 max pooling between layers. The classifications produced by this network were later cross-referenced using an algorithm described below.

We also build a variant of our two-stage postural tracking approach described above in order to identify when a bird was carrying a seed in its beak. This algorithm used the same first stage to coarsely track the bird’s head, body and tail, and to triangulate 3D position. However rather than cropping around the body, for seed carrying detection we cropped a tighter region centered on the bird’s face. A custom network implemented in TensorFlow 2^4^ was then used to predict for each frame whether the bird currently held a seed in its beak. This network consisted of 25 7×7×1 convolutions with stride 2, 50 3×3×25 convolutions with stride 1, 2×2 maxpooling, 50 3×3×50 convolutions, 100 3×3×50 convolutions, 2×2 maxpooling, 100 3×3×100 convolutions, global average pooling, 20% dropout, and a single linear output. All layers used SELU activations. This network was applied to the image acquired by each camera independently, and then outputs were summed across views and passed through a sigmoid nonlinearity to predict the probability of a bird carrying a seed for each frame. A hierarchical network combining simultaneous information from all 6 views was then trained end-to-end on manually annotated images.

The outputs of both the bottom camera and the seed carrying network predictions were then input to a user GUI used to annotate all of a bird’s cache site interactions in a semi-automated manner. We used a heuristic algorithm, described below, to identify possible interactions involving a cache or retrieval. We then generated flags requiring manual user review whenever our two independent algorithms were in disagreement. The first algorithm detected all times when the bird began or finished carrying a seed in its beak. If a bird gained a seed during a site manipulation, the site manipulation was proposed as a retrieval, and if a bird lost a seed it was carrying during a site manipulation, the action was proposed as a cache. The cumulative number of seeds currently in each site was then computed for the entire session. A second algorithm used bottom camera data, and made a prediction about whether a cache site contained a seed immediately prior to and subsequent to the site interaction. A flag was generated if the second algorithm detected an occupied site which the first algorithm predicted as empty, and vice-versa. A flag was also generated if a retrieval was detected from an empty site. The GUI allowed a manual annotator to browse through all flags, viewing the full sequence of all site manipulations and predicted seed contents at a site within that session, as well as behavioral and bottom camera video for each site manipulation. After manual correction by the annotator, the flag detection algorithm was re-run to ensure consistency of all bottom camera and seed carrying predictions with the updated annotations.

### Design of the electrophysiology implant

We designed a light-weight implant for electrophysiological recordings during behavior. The implant was designed for use with a 64-channel silicon probe (H6 ASSY-236, Cambridge NeuroTech), glued to an aluminum drive (nanodrive, Cambridge NeuroTech). The probe was connected to a custom built headstage that used a 64-channel amplifier (RHD2164, Intan Technologies). The headstage communicated digitally with a recording system (C3100 RHD USB interface board, Intan Technologies) over a digital SPI connection (C3216, RHD ultra-thin SPI interface cable, Intan Technologies) connected to a motorized commutator (Assisted Electrical Rotary Joint 24_PZN12, Doric Lenses). To minimize the forces exerted by the cable, strands of a thin elastic string (1 mm Flat Electric Crystal Stretch String) were tied to the cable to provide a low spring-constant force. The probe and drive were designed to fit within a 3D printed protective housing (Clear Resin, Formlabs), which consisted of two components. The top component housed the headstage, which formed its rear wall, as well as the nanodrive and the probe. The bottom component (base unit) was a small part that attached to the skull. The entire assembly was 1.2 g (0.1 g probe, 0.46 g headstage and connectors, 0.28 g nanodrive, 0.35 g housing).

### Surgical approach

Our surgical approach consisted of two steps, detailed below. In the first step, the implant site on the brain was prepared, and base unit of the implant was attached to the skull. In the second step, the top component of the implant was attached.

For the first step, chickadees were anesthetized using 1.5% isoflurane in oxygen. An injection of dexamethasone was made intraperitoneally (2 mg/kg). Fluids (0.9% NaCl, 0.1 mL every 45 min) were administered subcutaneously for the duration of the surgery. Feathers were removed from the top of the head around the surgical site, and the surgical site was cleaned using betadine and 70% ethanol solution. The chickadee was then placed in a stereotaxic apparatus, secured by custom designed ear bars and beak clamp. The bird’s head was aligned to stereotaxic axes by adjusting the beak clamp to 30° below horizontal. A silver ground wire (uncoated, 0.005” diameter) was inserted beneath the skull ∼1 mm anterior and 2 mm lateral to lambda, over the right hemisphere. A craniotomy and durotomy were then performed covering a 1×1 mm area centered 3 mm anterior to lambda and 0.6 mm lateral to the midline, over the left hippocampus. A 3D printed biocompatible resin insert, consisting of a 0.2 mm depth, 1 × 1 mm square underneath a 0.3 mm depth, 1.5 × 1.5 mm square, was inserted into the craniotomy site and cemented to bone. The insert contained a small central slit (0.4 × 0.1 mm) through which silicon probes could be later inserted. After the insert was cemented (RelyX Unicem, 3M), the space above the craniotomy was filled with a protective layer of Kwik-Cast (World Precision Instruments). The 3D printed base unit (Clear Resin, FormLabs) was then cemented into place, centered above the craniotomy and secured to the skull. A removable 3D printed cap was attached to the base unit to protect the craniotomy site. Buprenorphine (0.05 mg/kg) was injected intraperitoneally and the bird was allowed to recover for 1-2 weeks after this initial surgery.

After birds recovered from the initial surgery, a second procedure followed to implant the top component of the device. Birds were anesthetized and given dexamethasone as above, the removable cap was removed from the base unit, and Kwik-Cast was removed to expose the craniotomy site and the insert. Silicone gel (Dow DOWSIL 3-4680) was added to fully cover the exposed brain, insert, and ∼1 mm of space above. The silicon probe and nanodrive assembly were then positioned to allow probes to advance through the insert’s slit into underlying brain, and the nanodrive was cemented to the skull. The silicon probe and ground wire were connected to the headstage, and the headstage was inserted into protective housing, which was cemented onto the base unit. Birds were given 1 week to recover from this implantation before continuing behavioral and electrophysiology experiments.

### Electrophysiology protocol and spiking data pre-processing

We observed that neural signals degraded rapidly when silicon probes were left in neural tissue between experimental sessions. This degradation included decreases in the numbers of units and amplitudes of spikes, as well as increases in electrode impedance. We therefore developed a “semi-acute” recording protocol. Approximately 15-30 min before recording on an experimental day, the implanted microdrive was used to lower silicon probes to the desired depth in hippocampus. Recording depth was varied across sessions in the same bird, with data reported in this manuscript pooled across depths up to 1.5 mm below brain surface. Immediately following an experiment, probes were fully retracted such that they rested above the brain, with their tips embedded in the silicone gel covering the brain. Despite making repeated recordings along a similar recording track, we observed only gradual signal degradation over weeks of recording. Probe impedances and background noise also remained low throughout experiments when following this protocol.

Electrophysiology data was bandpass-filtered between 1 Hz and 10 kHz before digitization at 30 kHz with 0.2 μV resolution. Acquisition software simultaneously recorded the digital trigger used to acquire each frame of the behavioral video, for posthoc alignment. At each time point, the median across all channels was subtracted from all signals, and spikes were then extracted by the Kilosort 2.0 algorithm^5^.

For all units extracted by Kilosort, we collected a number of spike metrics used for both quality control and classification of excitatory and inhibitory units. First we obtained the average spike waveform, and from it calculated the spike width, asymmetry, the ratio of the peak and trough of the spike, and the ratio of the peak and trough of the waveform derivative. We additionally calculated the mean rate and a burst-index given by the ratio of the inverse median interspike interval and mean rate. Some of these features have been previously used to distinguish putative excitatory and inhibitory neurons in the avian hippocampus^6^. For quality control, we calculated the spatial extent of each unit along the probe, as well as the cluster contamination rate determined by Kilosort. All units were visualized using a tSNE embedding of these features, and manual clusters were identified for artifacts and neural spikes. A Gaussian mixture model was fit to these manually determined clusters and used to classify all units using the metrics described above. Artifacts had large spatial spread, high contamination rates, and/or unusual waveform shapes compared to neurons. Neurons were subdivided into three groups. Inhibitory neurons had higher and less bursty rates, with narrower and more symmetric spikes relative to excitatory neurons. Some neurons had properties intermediate between clearly excitatory and inhibitory clusters, and were labeled as unclassified neurons.

Spike times were aligned to behavioral videos, and spike counts were binned at 60 Hz. For analysis of single units, we considered only units classified as excitatory or inhibitory neurons, with contamination rates <0.2, and mean firing rates above 0.02 Hz. For population analyses, we included unclassified neurons, and neural units with contamination rates up to 1. We normalized each unit’s firing rate for population analyses by first subtracting its baseline rate, computed as a running 30 min average. After baseline subtraction, we divided firing rate by its standard deviation, which was regularized by adding a small number (0.6 Hz).

### Single-neuron place and cache tuning

Place maps were obtained by smoothing visit data to each perch with a LOESS quadratic model (MATLAB function *fit* type *loess*, with span 0.4). To quantify the significance of place tuning, we generated 1000 shuffle samples by circularly permuting neural data relative to the sequence of visit locations. For both the real data and the shuffled samples, we computed the standard spatial information index^7^. A cell was considered to be a place cell if the spatial information for the real data exceeded 95% of the shuffles.

To quantify the sparsity of cache responses, we calculated the fraction of caches for which a neuron’s firing rate was above its firing rate averaged across the entire session. For most excitatory neurons, which had average rates <1 Hz, this threshold was surpassed if any spikes occurred during a cache. Again, we compared this fraction to 1000 shuffled data points. In this case the shuffle distribution was generated by circularly permuting cache times relative to neural spike times. We used the 5^th^ and 95^th^ percentiles of the shuffled distribution to classify neurons as significantly suppressed or enhanced.

### Population vector correlation-based analyses

Most analyses of barcodes and reactivation used a similar analysis framework based on correlating pairs of neural population vectors for two events in the same session. For all analyses, before calculating correlations, the mean across all instances of an action (i.e. across all locations) was subtracted from each instance. For example, the mean across all caching events in a session was subtracted from each caching event. Pairs of events separated by less than 1 minute were excluded from the analysis. To obtain standard errors, the analysis was repeated 100 times while resampling with replacement the 54 sessions included in the full dataset. An exponential function with independent parameters for baseline, tau, and amplitude were fit to the data, excluding correlations between events at the same site involving any non-visit action. For analyses involving linear mixed-effects modeling, we used *fitlme* (MATLAB, 2022b) to fit the model and perform t-tests on the significance of each fixed effect.

The robustness of spatial coding is typically assessed using data averaged over long periods of time – e.g. correlating spatial maps from one half of a behavioral session to another^6, 8^. For typical session durations and smoothing parameters, these kinds of analysis in the hippocampus have produced average correlation values of ∼0.5. Repeating these analyses for our data set produced similar values. However, analysis reported in this paper is unusual in that it samples spiking during very brief single events, lasting ∼1 s. As expected from the variability of single trial spike counts in brief windows, the resulting correlation values were much smaller (0.024 for the average visit-visit correlation across the data set). All population vector correlations in the manuscript were multiplied by a constant factor (40.95) such that the value of the exponential curve for visit-visit correlations at the same site was exactly 1.

### Place code subtraction

We developed a method to estimate and remove the place code from caching data, in order to isolate the barcode component of activity during caching. For each site in the arena, we first estimated the place activity of each cell, by holding out that site and smoothing data from all perch arrivals at other sites using a LOESS quadratic model (MATLAB function *fit* type *loess*, with span 0.4). For this estimation, we included data from all perch arrivals including both those with and without site interactions. This procedure thus uses data from all other locations in the arena to estimate the smooth spatial component of a neuron’s activity, while excluding any contribution from site-specific activity during actions at a particular site. Place activity values across all neurons were combined into a “place vector” defined at each site. The mean value for each neuron across sites was subtracted, similarly to the way cache activity had been mean-subtracted, and place vectors were normalized to unit length. Finally, we computed the projection of cache activity onto this unit vector and subtracted it from the cache activity. This procedure avoids assumptions about how consistent the strength of place coding was across caches, sessions, and birds. “Barcode” activity in this paper refers to activity during a cache after this place subtraction procedure.

### 1D behavior on a circular track

For 1D behavior, an arena was constructed similarly to the 2D arena described above, but with only 20 perches arranged in a circle of 49.1 cm diameter. Perches were oriented tangentially to the circle. The arena was positioned in a square 60 × 60 cm enclosure. Motorized feeders (3D printed, white resin) were placed in the corners of this enclosure, on the outside of the circular track next to four of the perches. On top of each feeder was a small well filled with water, which was available at all four feeders even when the feeders were closed. Cache sites were placed on the outside of the track next to the other 16 perches, centered at 52.8 cm diameter. This dimension meant that the midpoints between the perches and sites were on a circle of circumference 160 cm; i.e., we considered 8 cm to be the arc distance between two adjacent sites. The cache sites used a previously published^1^, version of the design.

Each bird was typically recorded in 12 sessions on separate days, with at least one day of rest between sessions. One of the feeders was chosen as the “rewarded” feeder for the first three sessions, then a different feeder was chosen for the next three sessions, and so on. For each bird, the four feeders were rewarded in a different random sequence. Each session started with the feeder closed for 10 min and ∼3 of the sites baited. The rewarded feeder was then opened for 5 min every 30 min for the duration of the 1.5-2 h session.

One camera (Edmund Optics, EO-2323C) was mounted on the ceiling of the enclosure and used to monitor the position of the bird. Do detect position, we used a neural network^9^ trained on the center of the bird’s head. A second, identical camera was mounted in the wall of the enclosure. A third camera (Edmund Optics, acA2500-60uc) was mounted under the floor of the arena and used to monitor the contents of the caches. Caches, retrievals, and checks in the 1D arena were detected using a semi-automated annotation procedure^1^.

### Calcium imaging

Experiments on the circular track were performed before our lab had developed technologies for silicon probe recordings. For these experiments, we used calcium imaging with head mounted microscopes. Calcium imaging did not allow some of the analyses that we performed in the 2D arena (e.g. analysis of inhibitory cells or the highly temporally precise analyses of neural dynamics). However, it provided comparable numbers of recorded cells per session and could be used for analyzing the spatial code – which was the main purpose of the circular track.

Surgery for calcium imaging consisted of injecting a virus containing GCaMP6f, as well as implanting a GRIN lens and baseplate, using procedures previously described^10^. Anesthesia, initial preparation of the bird, and analgesia were done as for silicon probe experiments described above. Six small holes were drilled through just the top layer of the skull, for the purpose of anchoring the implant. The main craniotomy was then made, centered at 3.25 mm anterior and 0.7 mm lateral to lambda. Before the craniotomy on the inner layer of skull, an antibiotic solution (Baytril 3.8 mg/mL) was applied to the surface of the skull for five minutes, then wicked away. Insect pins were inserted through neighboring pairs of the anchor holes, and both the insect pins and the anchor holes were covered with cement (D69-0047, Pearson Dental). The craniotomy and durotomy were then completed.

Birds were injected with the AAV9-CAG-GCaMP6f-WPRE-SV40 virus (100836-AAV9, Addgene). The total amount of virus injected was 897 nL (65 injections of 13.8 nL each), using a Nanoject II (Drummond Scientific) with a pulled glass pipet tip. Throughout the injection, the surface of the brain was covered in Kwik-Sil (World Precision Instruments). There was a 10 sec waiting period between injections, and a 25 min waiting period after the final injection, before withdrawing the pipet and removing the Kwik-Sil.

Before implanting the GRIN lens, the head was angled into a typical chickadee resting head posture (beak bar approximately 10 degrees below the horizontal). A GRIN lens (1 mm diameter, 4 mm length, 1050-004595, Inscopix) was implanted directly over the injection site, any remaining exposed brain was covered with Kwik-Sil, and the lens was anchored in place using cement (D69-0047, Pearson Dental). Next, a baseplate was positioned into a good focal plane with the Inscopix miniscope (focusing slightly below the brain surface, with the miniscope objective approximately 300 µm above the surface of the GRIN lens), and the baseplate was cemented in place. The surface of the cement was covered with black nail polish to prevent light contamination.

Inscopix microscopes have a heavier cable than our silicon probe implants. To support the weight of the cable, birds were therefore fit with a leg-loop harness^11^ two weeks after surgery. The harness remained permanently on the bird and consisted of a 3D-printed plastic attachment for the cable (20 × 10 ×6 mm) pressed against the bird’s back and two loops of elastic string (39 mm for each leg, Outus Elastic Cord) that were hooked onto the bird’s thighs. During behavioral sessions, the microscope was snapped into the magnetic headplate, and the cable at a point ∼13 cm from the microscope was attached to the harness.

### Analysis of calcium imaging data

For analysis of calcium imaging data, we use procedures previously described^10^. Imaging data were collected at 20 fps. Neuronal traces were extracted from raw fluorescence movies using a constrained non-negative matrix factorization algorithm intended for 1-photon calcium imaging data (CNMF_E^12, 13^). We used a multi-scale approach^14^ to extract stable fluorescent traces from long videos (2 h in our case). Before applying CNMF_E to the raw videos, we applied a motion correction algorithm^15^. The vast majority of data contained no motion above 1 pixel RMS shift.

The multi-scale CNMF_E approach was run in three steps. First, data were averaged in bins of 20 frames, then temporally downsampled by a factor of 20. Cell footprints were found in the downsampled movie using the standard CNMF_E algorithm. These footprints were then used to extract temporal traces on segments of the non-downsampled data. Finally, the raw traces were deconvolved to detect the time and amplitude of each calcium event. To eliminate some infrequent imaging artifacts, any calcium events with an amplitude greater than 1.5 times larger than the 99^th^ percentile of all calcium events for that cell were eliminated from all analyses. For all firing rate calculations in the paper, events were weighed by their amplitude.

### Effects of caching on the place code

To determine the effect of caching on spatial tuning, we used data from the circular track described above. For most analyses, the bird’s position was classified as being at one of 20 sites if the radius from the center of the arena was between 19.75 and 30 cm, and the angle was within ±9° from the center of the perch of the site. For the distribution of place fields shown in Figure 4, the circular track was instead divided into 60 segments 6° wide, with every third segment centered on a perch.

For each cache and each cell, we first calculated the spatial tunings of the cell before and after the cache. For pre-cache tuning, we used the time period starting at the last cache or retrieval that previously occurred at the same site; if no such event occurred, the period started at the beginning of the session. The period ended 5 min before the cache. For post-cache tuning, the period started 5 min after the cache and ended at the next cache or retrieval, if such existed, or at the end of the session. Periods of 5 min before and after the cache were excluded because behavior during these periods was typically very different from other parts of the session when the bird moved mostly consistently around the circle. We also excluded periods of ±500 ms around checks during both the pre- and the post-cache periods. For both periods, we determined the number of calcium transients and the behavioral occupancy at each of the 20 sites in the arena. If any site had <1 s occupancy either before or after the cache, the cache was excluded from the analysis. Calcium transient counts and occupancy were each smoothed by a 3-point square window, and firing rates were calculated by dividing smoothed counts by smoothed occupancy values.

We compared pre-cache and post-cache tuning using four metrics. The first metric was the relative difference in peak firing rates, measured as the absolute difference between pre and post values, divided by the average of those values. For this analysis, peak firing rate was defined as the maximum of the 20 values across sites. For the remaining metrics, we only included cells whose peak firing rate was >1 event/s in both the pre and post periods. The second metric was the absolute difference in the positions of the peaks in spatial tuning, measured as the length of the shortest arc around the circular track between these positions. The third metric was the relative difference in the widths of the tuning curves, again measured as the absolute difference divided by the mean. Width was defined at half of the maximum of the tuning curve. The fourth metric was the cross-correlation of the pre and post tuning curves.

To compute statistical significance of each of the four metrics, we generated 100 shuffle samples in which the entire set of times included in the pre and post periods was shifted in time by a random amount, but ensuring that the entire pre and post periods continued to overlap with the session. The 99% confidence intervals for each of the metrics were then computed as 2.8 standard deviations of this shuffle distribution.

### Seed tuning

We determined the significance of the modulation of each neuron’s activity during checks by the occupancy of the site. For each neuron, we generated single trial responses by calculating firing rate in the window from 100 to 1000 ms from each check onset, and subtracting the baseline rate in the 1000 ms before check onset. These difference were then separately averaged across all checks of empty sites and all checks of sites occupied by a seed. Seed tuning was calculated as the difference between the average occupied and empty responses. To determine significance of this tuning, we generated 1000 shuffle samples by circularly permuting site occupancy assignment (occupied or empty) with respect to neural data during the checks.

### Population projections of place code, barcode, and seed code

To obtain the time course of population coding for place, barcode, and seed components, we defined three population vectors. The population vector for seed coding was defined as the difference in check responses for occupied versus empty sites, using the procedure described above, measured across all neurons. The place code vector was defined as that obtained using the leave-one-out procedure described above for place code subtraction. The barcode vector was defined as the average barcode activity (i.e. activity during caches after place code subtraction) across all caches at a site. All projections were cross-validated by recomputing vectors for each event while holding out the data to be projected. Standard errors were computed by 1000 bootstrap samples, drawing with replacement from the 54 session-averaged projection time courses.

---

**Supplementary Figure 1.**
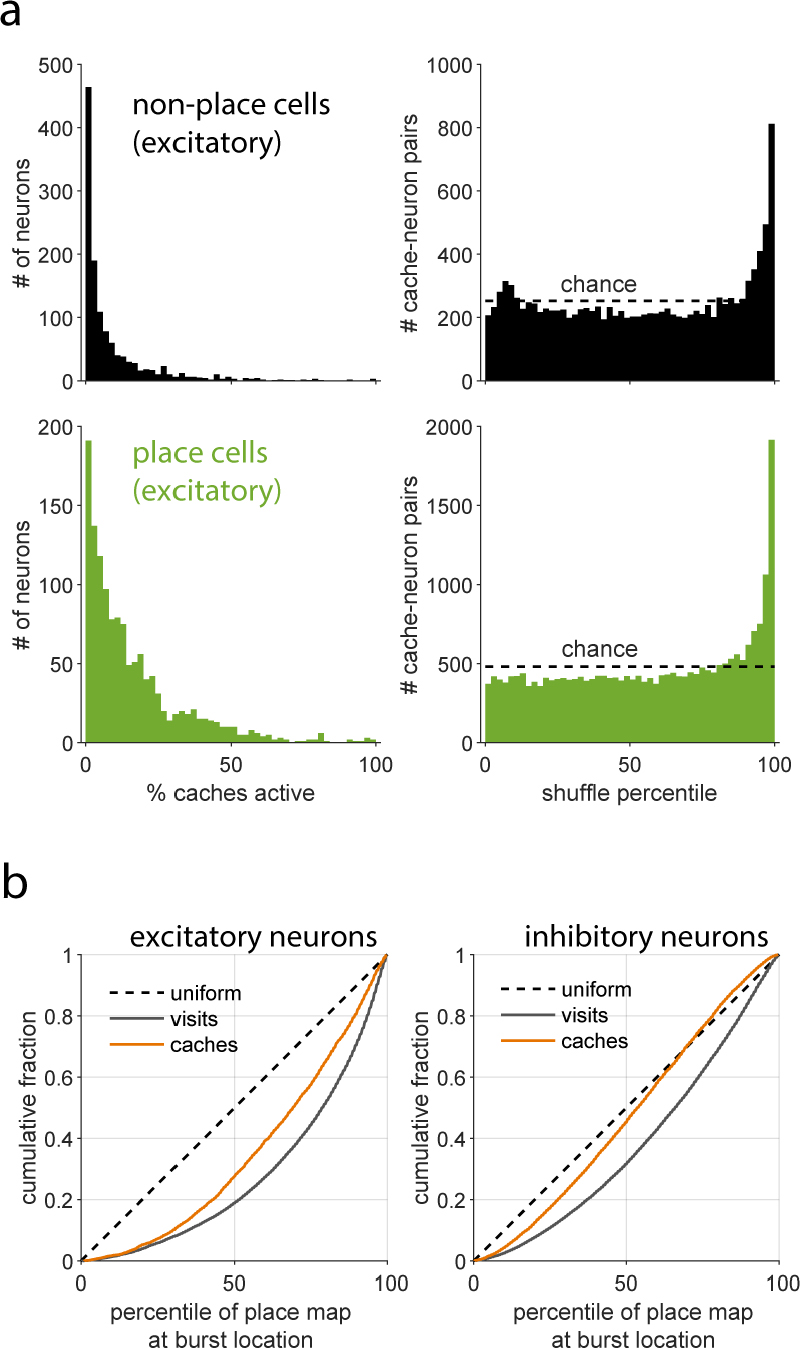
Cache responses in place and non-place cells. **a)** Analysis of excitatory neuron activity during caching as in Fig. 2b-c, but separately for neurons without (top) or with (bottom) significant place tuning. Left: fraction of caches on which a neuron responded (i.e. exceeded mean firing rate). Right: Comparison of cache responses to shuffle distribution. Both place and non-place cells were sparsely active during caches, although place cells were less sparse, presumably because they exhibited place activity during caches within their place fields. Both place and non-place cells exhibited large bursts of activity during caches that were in the 95-100^th^ percentile of their shuffle distributions, meaning they were greater than the neuron’s rate at almost any other time. Thus place and non-place cells exhibited similar bursts of activity during caching. **b)** Analysis of the relationship between cache response locations and a neuron’s place map. For each neuron, we determined the location of all cache responses in the 95-100^th^ percentile of their shuffle distribution, and calculated the percentile of the neuron’s place map at each of these location. The cumulative histogram of these percentiles is plotted in orange. This analysis was repeated on data during visits for comparison (grey). If cache responses were distributed uniformly throughout the arena, irrespective of a neuron’s place activity, the cumulative histogram would be a straight line (dashed black line). Analysis was repeated for excitatory (left) and inhibitory (right) neurons. Cache response were more clustered around a neuron’s place field than chance, but were more dispersed than activity during visits. This is consistent with cache responses being driven by a combination of place tuning and a caching-related, site-specific component (i.e. the barcodes).

**Supplementary Figure 2.**
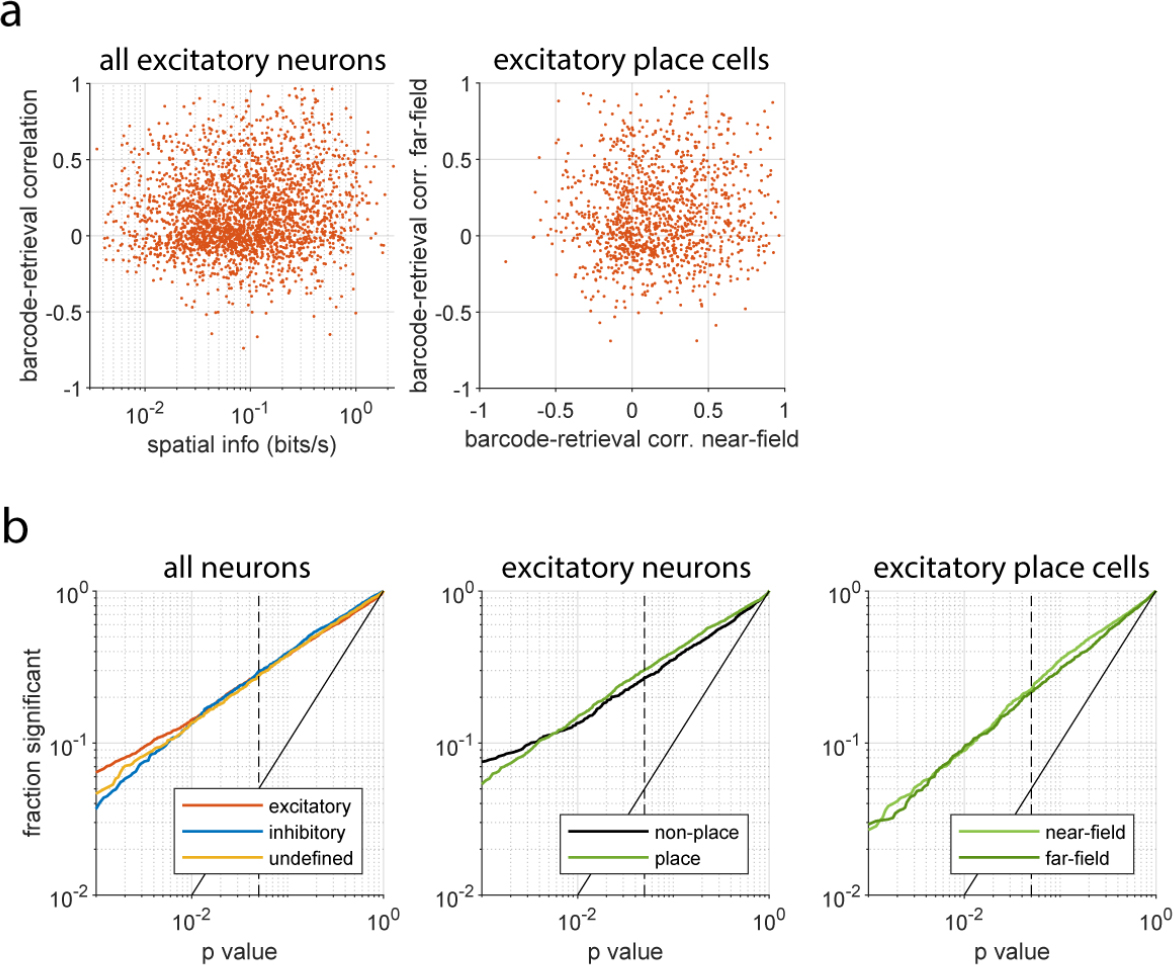
Barcodes are mostly unrelated to place tuning. **a)** Left: the spatial information of a neuron’s place map plotted against single-neuron barcode-retrieval correlation. Single-neuron barcode-retrieval correlations were calculated by computing barcode activity as described in the main text, and calculating a vector for each neuron of its barcode activity across all sites (averaging across multiple caches at a site when applicable). A vector was similarly calculated for retrievals, and a neuron’s barcode and retrieval activity was then correlated across all sites with at least one cache and one retrieval. The relationship between the strength of a neuron’s place tuning and the strength of its barcode tuning was significant, but very weak (spearman correlation: p < 0.001, r = 0.11). Right: in order to determine whether there was a relationship between a neuron’s place field and its barcode activity, we recomputed the barcode-retrieval correlation for each place cell after dividing all sites in the arena into non-overlapping halves that were closer or further from the peak of the neuron’s place field (‘near-field’ and ‘far-field’). Place cells participated heterogeneously in barcodes for caches near and far from their place field, although there was a small but significant difference in mean correlation (near: 0.17, far: 0.14, p < 0.01, t-test). **b)** The fraction of neurons with significant barcode-retrieval correlations plotted for a range of p values. Significance was determined by randomly shuffling the site locations of the barcode and retrieval vectors. A solid black diagonal line shows the expected fraction of significant neurons for each p value threshold, and a dashed vertical line indicates a p value of 0.05. Left: barcode significance for all neurons grouped by neuron type; middle: for all excitatory neurons grouped by significance of place tuning; right: for excitatory place cells using correlations computed only at sites near or far from the neuron’s place field. Note the slight decrease in significant fraction for rightmost plot is due to significance being tested on two separate halves of the data used in left and center plots. Barcode tuning had similar significance across cell types, for place- and non-place cells, and at different distances from place fields.

**Supplementary Figure 3.**
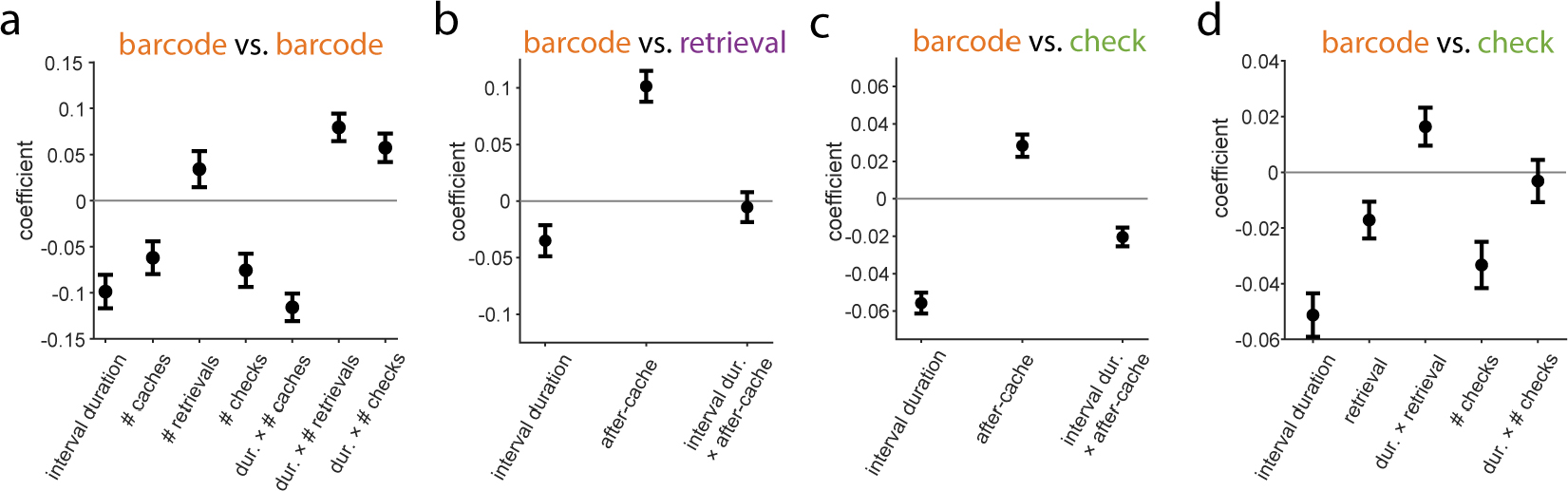
Barcode reactivation reflects experience at a cache site. **a)** Coefficients of a linear mixed-effects model of the correlation between two cache barcodes at the same site. The model contains random effects for each session (not shown) and fixed effects for the duration of time between caches, the number of intervening caches, retrievals, and checks, and nonlinear interactions between duration and each intervening event count. Coefficients are without units, since variables were z-scored before model fitting. All coefficients are significantly different from 0 with p < 0.001 except the linear term for # retrievals p = 0.082. Model coefficients are consistent with effects shown in Fig. 4a-b. Specifically, barcode correlations are reduced by both the number of intervening caches and the time between caches. The strongest coefficient was a negative interaction between duration and intervening caches, meaning that decorrelation between barcodes was particularly strong when caches were separated by both a long interval and multiple intervening caches. **b)** Coefficients of a linear mixed-effects model of the correlation between cache barcodes and retrievals at the same site. The models contained random effects for each session, and fixed effects for the duration between events, whether the retrieval was after or before the cache (binary variable), and a nonlinear interaction between duration and this binary variable. To match Fig. 4d, only barcode-retrieval pairs with no intervening caches or retrievals were analyzed. Model coefficients are consistent with results shown in Fig. 4d. Barcode-retrieval correlations were greater for retrievals after a cache than before a cache (p < 0.001), and decreased for longer intervals (p = 0.011). The interaction term was negligible (p = 0.69), implying that the post-cache increase in correlation was temporally stable. **c)** As in b), but for barcode-check pairs. All coefficients were significantly different from 0 with p<0.001. Model coefficients are consistent with results shown in Fig. 4f. Specifically, barcode-check correlations increased following a cache, and the negative interaction indicates this effect decayed with greater duration. **d)** Coefficients of a linear mixed-effects model of the correlation between cache barcodes and checks at the same site, only for pairs where the cache preceded the check, and with no intervening caches. The model contains random effects for each session, and fixed effects for temporal duration, whether an intervening retrieval had occurred (binary variable), the number of intervening checks, and nonlinear interactions between duration and the other two variables. Model coefficients are consistent with effects shown in Fig. 4f. The negative coefficient for retrieval (p < 0.01) and its positive interaction with duration (p = 0.016) implies that barcode-check correlations decreased after the cache was retrieved, particularly if the duration between cache and check was brief. We additionally observed effects of duration and intervening checks (p <0.001) with an insignificant interaction (p = 0.68). Error bars in all panels: s.e.m.

**Supplementary Figure 4.**
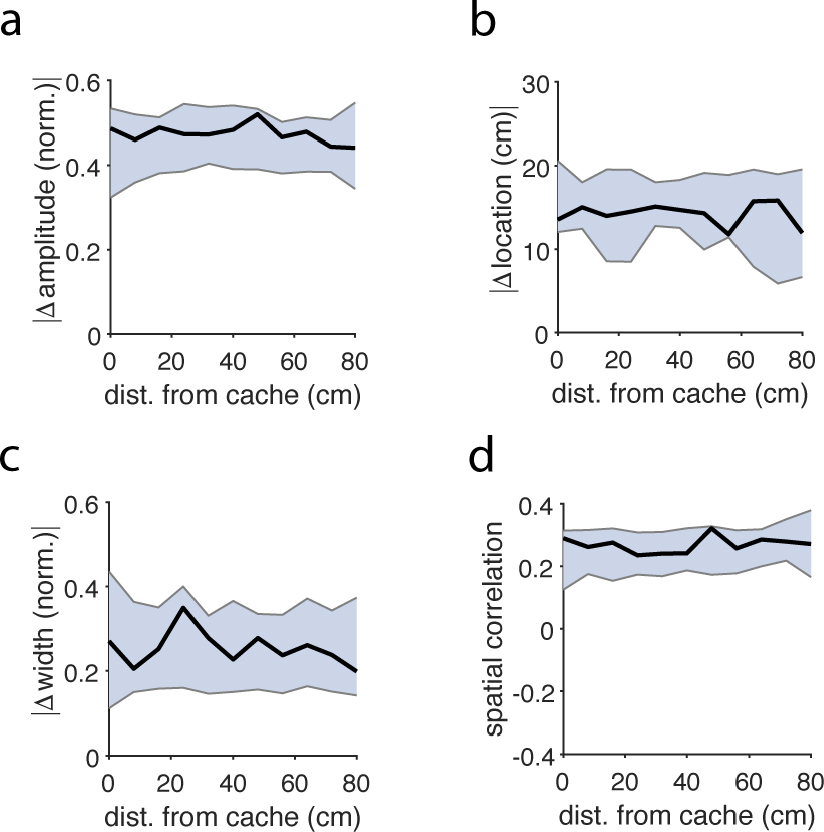
Place fields are unchanged following caches. **a)** The change in amplitude of a neuron’s place field, measured as the absolute difference in amplitude before and after a cache, normalized by the average of these two values. Mean change is plotted as a function of distance from the cache location to the peak of a neuron’s place field. Shaded area is the 99% confidence interval of a shuffle using randomized cache times. **b**) As in a) but for change in place field location. **c)** As in a) but for change in place field width. **d)** As in a) but for spatial correlation of a neuron’s place tuning before and after a cache. For all panels, changes to place tuning aligned to caches were similar to changes aligned to random, non-caching time points in the session. Caches thus did not lead to significant changes in place tuning in our experimental conditions.

**Supplementary Figure 5.**
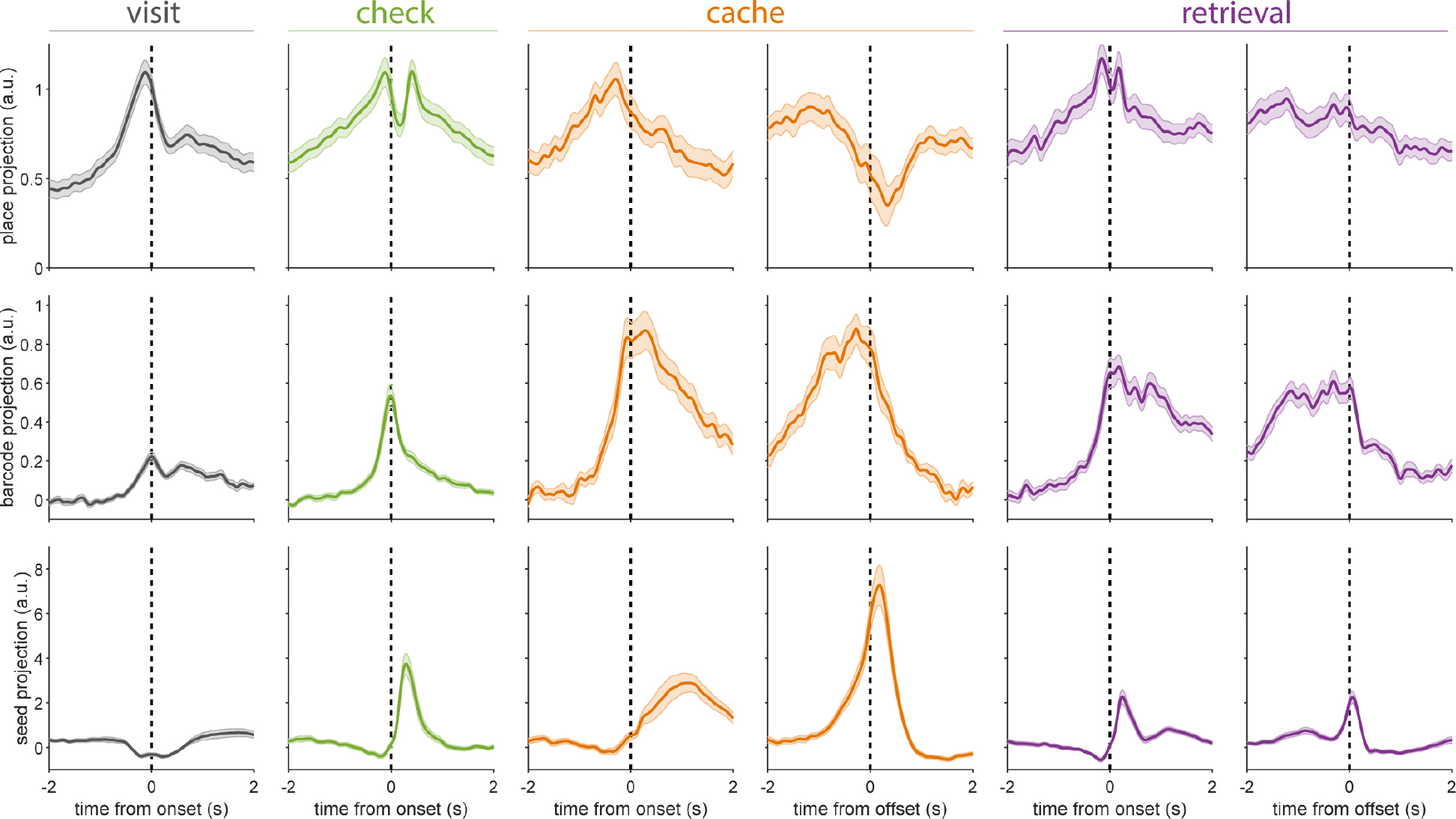
Temporal dynamics of place, barcode, and seed tuning. As in Fig. 5, projection of neural activity onto dimensions of place, barcode, and seed tuning. Projections are plotted for visits, checks, caches and retrievals aligned to onset, and for caches and retrievals aligned to offset. Projection values were scaled by a constant across all plots such that the place projection at time 0 was equal to 1. For seed projection plots, only occupied visits and checks are shown (see Fig. 5 for empty visits and checks). Shaded region is s.e.m. with bootstrap resampling, n = 54 sessions. The temporal sequence of place, barcode, and seed tuning during checks resembled a compressed version of that seen during caches, perhaps reflecting their briefer duration.

**Supplementary Movie 1. Example of caching behavior**

Example of caching behavior in one bird that was later implanted and recorded from. Initially, full-frame video data from one camera is shown at 1× real speed. The video is cropped, zoomed, and slowed to 1/8x speed when the bird makes a cache to better demonstrate usage of the cache site. Another camera’s view of cache site contents through the transparent bottom is overlaid during this time. Playback is restored to 1× speed after the first cache.

**Supplementary Movie 2. Example checking behavior with 3D postural tracking**

Example of checking behavior with an overlay of 3D postural tracking data. The video is played at 1/2x real speed. 3D postural keypoints are reprojected onto video data from all 6 behavioral videos and connected by a ‘skeleton’. Keypoints on the bird’s left, midline, and right side are plotted in yellow, green, and blue shades. Raw video data from each camera view is cropped and zoomed to simulate translational camera movement, keeping the bird centered and at constant physical distance, based on an estimate of the bird’s 3D position. At the start of the video the bird finishes caching a seed, and then makes a series of brief checks of cache sites as it moves across the arena.

